# Stiffness Regulates Intestinal Stem Cell Fate

**DOI:** 10.1101/2021.03.15.435410

**Authors:** Shijie He, Peng Lei, Wenying Kang, Priscilla Cheung, Tao Xu, Miyeko Mana, Chan Young Park, Hongyan Wang, Shinya Imada, Jacquelyn O. Russell, Jianxun Wang, Ruizhi Wang, Ziheng Zhou, Kashish Chetal, Eric Stas, Vidisha Mohad, Marianna Halasi, Peter Bruun-Rasmussen, Ruslan I. Sadreyev, Irit Adini, Richard A. Hodin, Yanhang Zhang, David T. Breault, Fernando D. Camargo, Ömer H. Yilmaz, Jeffrey J. Fredberg, Nima Saeidi

## Abstract

Does fibrotic gut stiffening caused by inflammatory bowel diseases (IBD) direct the fate of intestinal stem cells (ISCs)? To address this question we first developed a novel long-term culture of quasi-3D gut organoids plated on hydrogel matrix of varying stiffness. Stiffening from 0.6kPa to 9.6kPa significantly reduces Lgr5^high^ ISCs and Ki67^+^ progenitor cells while promoting their differentiation towards goblet cells. These stiffness-driven events are attributable to YAP nuclear translocation. Matrix stiffening also extends the expression of the stemness marker Olfactomedin 4 (Olfm4) into villus-like regions, mediated by cytoplasmic YAP. We next used single-cell RNA sequencing to generate for the first time the stiffness-regulated transcriptional signatures of ISCs and their differentiated counterparts. These signatures confirm the impact of stiffening on ISC fate and additionally suggest a stiffening-induced switch in metabolic phenotype, from oxidative phosphorylation to glycolysis. Finally, we used colon samples from IBD patients as well as chronic colitis murine models to confirm the *in vivo* stiffening-induced epithelial deterioration similar to that observed *in vitro*. Together, these results demonstrate stiffness-dependent ISC reprograming wherein YAP nuclear translocation diminishes ISCs and Ki67^+^ progenitors and drives their differentiation towards goblet cells, suggesting stiffening as potential target to mitigate gut epithelial deterioration during IBD.

---

Upon migrating on the soft basement matrix (BM) from the bottom of the crypt to the tip of the villus, intestinal stem cells (ISCs) differentiate to diverse types of gut epithelial cells, including Paneth cells, goblet cells, enteroendocrine cells (EECs), tuft cells, microfold (M) cells and enterocytes ^1^. Inflammatory bowel disease (IBD), which includes ulcerative colitis (UC) and Crohn’s disease (CD), is associated with the deterioration of gut epithelium, including reduction of ISCs ^2^ and increase of M cells in UC ^3^. Furthermore, due to the excessive secretion of collagen, the BM stiffens ^4–6^. It has been demonstrated that the stiffness of the BM can regulate the differentiation of mesenchymal stem cells, the progenitor cells of central nervous system and pancreatic progenitors ^7–9^. Yet, it is unclear how the BM stiffening in IBD impacts the fate of ISCs and their differentiation, and contributes to the epithelium deterioration.

## Quasi-3D gut organoids cultured on soft hydrogel matrix

To investigate the impact of BM stiffening on the differentiation of ISCs, we developed a platform for culturing quasi-3D gut organoids on top of soft polyacrylamide-hydrogel matrix (Fig. 1A). ISCs and their crypts were harvested from mice and seeded on the hydrogel matrix. *Lgr5-EGFPIRES-creERT2* mice were used to track Lgr5^+^ ISCs (Extended Data Fig. 1). As the organoids grew, the soft hydrogel surface buckled (0.6 kPa, matching that of a healthy BM ^6^), forming a quasi-3D structure that mimicked the invagination of the *in vivo* crypts (Fig. 1A, and 3D confocal imaging in Fig. 1L). The crypt-like regions were densely populated by the ISCs intermixed with the large, optically dark, UEA^+^ Paneth cells (Fig. 1B). The peripheries of the crypts were surrounded by the transit-amplifying (TA) progenitor cells with strong Ki67 expression, and Ki67 was also weakly expressed in Lgr5^high^ ISCs (Fig. 1B and Extended Data Fig. 2A). The villus-like regions were populated by Villin^+^ enterocytes (Fig. 1B), Muc2^+^ goblet cells, and Chromogranin-A^+^ (Chro-A) EECs (Extended Data Fig. 2B). Notably, the villus-like regions also exhibited a turnover rate of approximately 3 days (Extended Data Fig. 3), similar to that observed *in vivo*. By culturing these quasi-3D gut organoids on the hydrogel matrix of varying stiffness, we analyzed the impact of stiffness on the fate of ISCs and their preference of differentiation directions.

**Figure 1.**
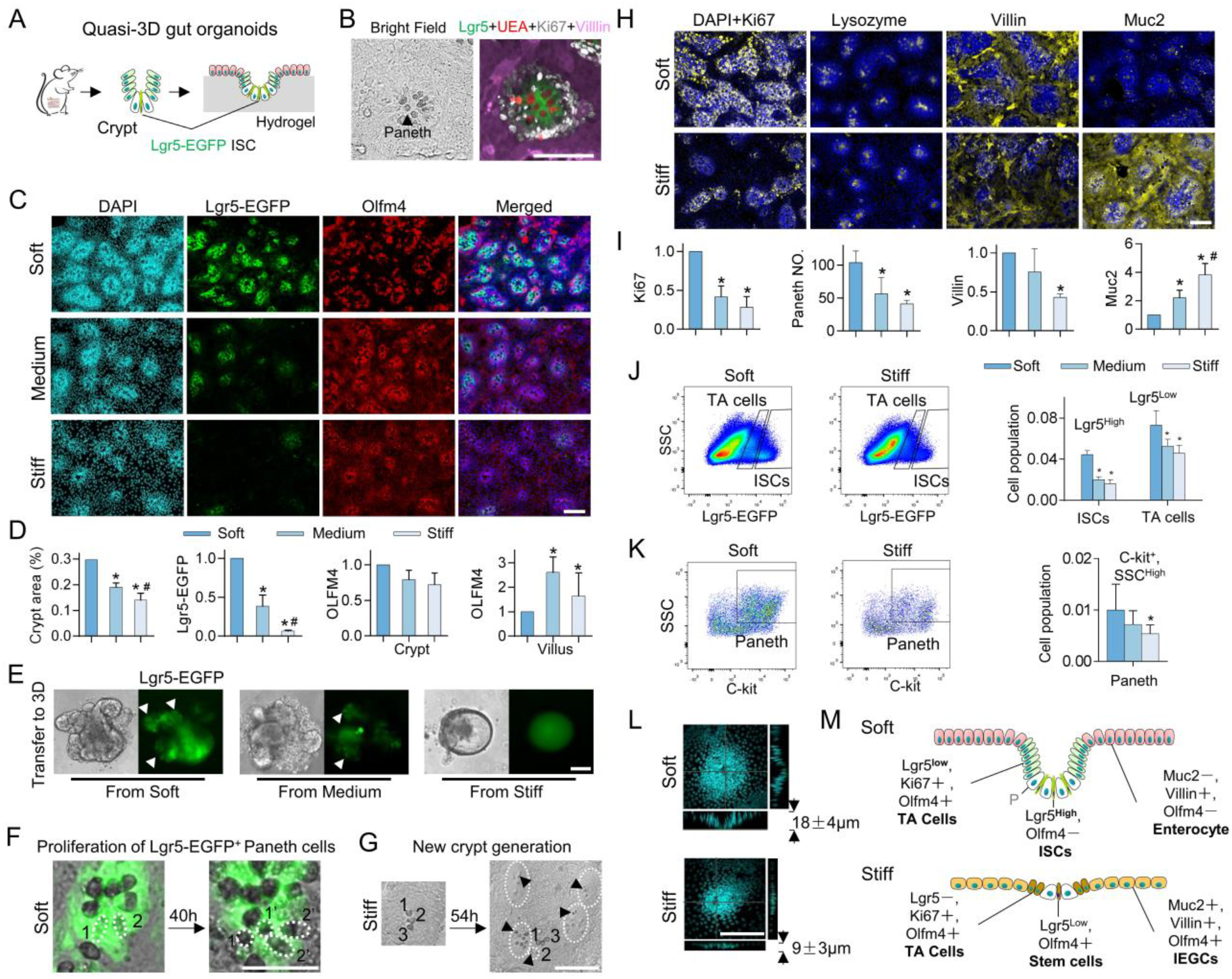
Stiffness determines the fate of ISCs. (A) Illustration of the experimental system. (B) Lgr5-EGFP^+^ ISCs were intermixed with the optically dark UEA^+^ Paneth cells, which were surrounded by Ki67^+^ TA cells in the crypt-like regions. The villus-like regions were populated by Villin^+^ differentiated cells. (C) The matrix stiffening from soft (0.6kPa) to medium (2.4kPa) to stiff (9.6kPa) reduced the size of the crypt-like regions with the dense nuclei and decreased the expression of Lgr5. Stiffening extended Olfm4 into the villus-like regions. (D) Quantification of the fluorescent intensity per unit area of crypt / villus regions. The crypt and villus regions were segmented using customized code based on DAPI intensity (Method, *n*=3-5). (E) The 3D organoids derived from the soft and medium matrix budded with Lgr5-EGFP^+^ ISCs (white arrows). The 3D organoids derived from the stiff matrix grew more like Lgr5-EGFP^−^ cysts. *n*=3. (F) Lgr5-EGFP^+^ ISCs (1 and 2) differentiated into two Paneth cells (1’ and 2’) on the soft matrix (Movie S1, *n*=3). (G) On the stiff matrix cells in the villus-like regions differentiated into Paneth cells (black arrows), which was followed by the new crypt generation (white dashed line, Movie S2, *n*=3). (H) The stiffening decreased the expression of Ki67, Lysozyme and Villin, but increased Muc2, as quantified via fluorescent intensity (I, *n*=3-5). (J and K) Flow cytometry analysis showed that stiffening decreased Lgr5^high^ ISCs, Lgr5^low^ TA cells, and Paneth cells (*n*=3). (L) 3D confocal imaging showed that the stiffening significantly inhibited the crypt invagination (*P*<0.05, *n*=3). (M) A schematic summarizes the impact of stiffening on all cell types. ‘P’, Panetch cell. Scale bar, 100 μm. * V.S. Soft and # V.S. Medium, *P*<0.05 (Student’s *t*-test).

## Stiffening reduces the number of Lgr5^+^ ISCs and promotes their differentiation into immature enterocyte-goblet cells (IEGCs)

Increasing the matrix stiffness from soft (0.6 kPa, matching that of a healthy BM ^6^) to medium (2.4 kPa) to stiff (9.6 kPa, matching that of an inflamed BM ^6^) gradually decreased the crypts surface area and reduced the number of Lgr5-EGFP^+^ ISCs (Fig. 1C and 10-days live-cell imaging for Lgr5-EGFP in Extended Data Fig. 4A). To further verify the impact of stiffness on ISCs, after 11 days of culture, the cells were detached from the hydrogel matrix and transferred to the inside of Matrigel^®^ to grow 3D organoids (Method). The 3D organoids from the soft or medium matrix budded to form the crypt regions with the Lgr5-EGFP^+^ ISCs, but those from the stiff matrix grew more like cysts with a significantly smaller number of buddings (Fig. 1E and quantified in Extended Data Fig. 4B), confirming the loss of stemness on the stiff matrix.

In addition to Lgr5, the expression and distribution of another stem cell marker, Olfactomedin 4 (Olfm4), exhibited strong correlation with the stiffness. On the soft matrix, Olfm4^+^ cells were concentrated in the periphery of the crypt-like regions (Fig. 1C and 1D). Upon increasing stiffness, Olfm4^+^ cells became interspersed throughout the crypt region and replaced the Lgr5^+^ ISCs to directly border with Paneth cells (Figs. 1C and Extended Data Fig. 5). Notably, on the stiff matrix, Olfm4 expression extended into the villus-like regions.

Live-cell imaging on the soft matrix showed that Lgr5-EGFP^+^ ISCs divided and differentiated into large and optically dark Paneth cells (Fig. 1F and Movie S1) which, in turn, contributed to the maintenance of ISC niche and stemness ^10^. In contrast, Lgr5-EGFP^+^ ISCs greatly diminished on the stiff matrix (Fig. 1C and Extended Data Fig. 4A). However, on the stiff matrix the cells in the villus-like regions differentiated into Paneth cells and ultimately generated ectopic, new crypt-like regions (Fig. 1G and Movie S2). The incidence of these ectopic crypts was approximately three-fold higher on the stiff matrix compared to the soft matrix (Extended Data Fig. 4C and Movie S3).

Stiffening also altered the proportion of differentiated cells. Stiffening diminished Ki67^+^ proliferating progenitor cells and Lysozyme^+^ Paneth cells, as well as the expression of the enterocyte markers, Villin and Alpi ^11^ (Fig. 1H and I, and Alpi in Extended Data Fig. 6). The EEC marker - Chro-A was also decreased by stiffening (Extended Data Fig. 6). In contrast, markers of secretory progenitor cells- Dll1 ^12^ and goblet cells- Muc2 were increased (Fig. 1H and I, and Dll1 in Extended Data Fig. 6). Notably, on the stiff matrix a new cell type emerged in the villus-like regions that co-expressed the markers for enterocytes, goblet cells, and stem cells (i.e., Villin, Muc2, and Olfm4, Fig. 1C, 1H and Extended Data Fig. 7). We named this new cell type the immature enterocyte-goblet cell (IEGC), where ‘immature’ refers to the co-expression of the different cell type markers. Flow cytometry analysis (Method) confirmed the reduction of ISCs (Lgr5-EGFP^high^), TA cells (Lgr5-EGFP^low^) and Paneth cells (CD24^high^, C-kit^+^ and SSC^high^) ^13, 14^ on the stiff matrix (Fig. 1J and K). Interestingly, 3D confocal imaging showed that the depth of the crypt-like regions was two-fold greater on the soft matrix compared to the stiff matrix (Fig. 1L).

The schematic in Fig. 1M summarizes the impact of stiffening on the various cell types: in the interior of crypt-like regions, stiffening led to the loss of Lgr5^high^ ISCs cells and the appearance of Olfm4^+^ stem cells which were directly adjacent to Paneth cells. It also reduced Lgr5^low^, Ki67^+^ TA progenitor cells. In the villus-like regions, stiffening led to the replacement of the Villin^+^ mature enterocytes by Muc2^+^, Villin^+^, and Olfm4^+^ IEGCs.

## The impacts of stiffness on ISC fate are YAP-dependent

Matrix stiffening stimulated YAP expression (Fig. 2A and 2D) and promoted YAP nuclear translocation (Fig. 2A and Extended Data Fig. 8). Lgr5^high^ ISCs were YAP negative, and YAP expression was inversely correlated with Lgr5 expression (Fig. 2A). To better distinguish the expression patterns of cytoplasmic YAP (cyto-YAP) and nuclear YAP (nuc-YAP), we performed immunostaining for non-phosphorylated YAP and the Ser 127 phosphorylated YAP, since YAP nuclear translocation is negatively regulated by YAP phosphorylation ^15^. The non-phosphorylated YAP expression was uniformly increased across both the crypt- and villus-like regions on stiff matrix and showed pronounced nuclear localization (referred as nuc-YAP for simplicity in Fig. 2B, quantified in 2D). In contrast, phosphorylated YAP that is primarily cytoplasmic exhibited strong region-dependent expression, i.e., it was decreased by the stiffening in the crypt-like regions, but increased in the villus-like regions (referred as cyto-YAP in Fig. 2C, quantified in 2D). To assess the relationship between YAP expression and the ISC differentiation trajectory, we counterstained total YAP with the markers of the differentiated cells. Proliferating cell marker (Ki67, Extended Data Fig. 9) and stem cell marker (Olfm4, Fig. 2E and crypt-like region in Extended Data Fig. 10) were both positively correlated with cyto-YAP, whereas goblet cell marker was positively correlated with nuc-YAP (Muc2, Fig. 2F). The Villin^+^ enterocytes on the soft matrix were YAP negative, and the Villin expression tended to decrease when YAP showed nuclear localization on the stiff matrix (Fig. 2G). Notably, the epidermal growth factor receptor (Egfr) which is involved in the process of inflammation and cancer ^16^ was positively correlated with cyto-YAP (Extended Data Fig. 11), similar to the stem cell marker, Olfm4.

**Figure 2.**
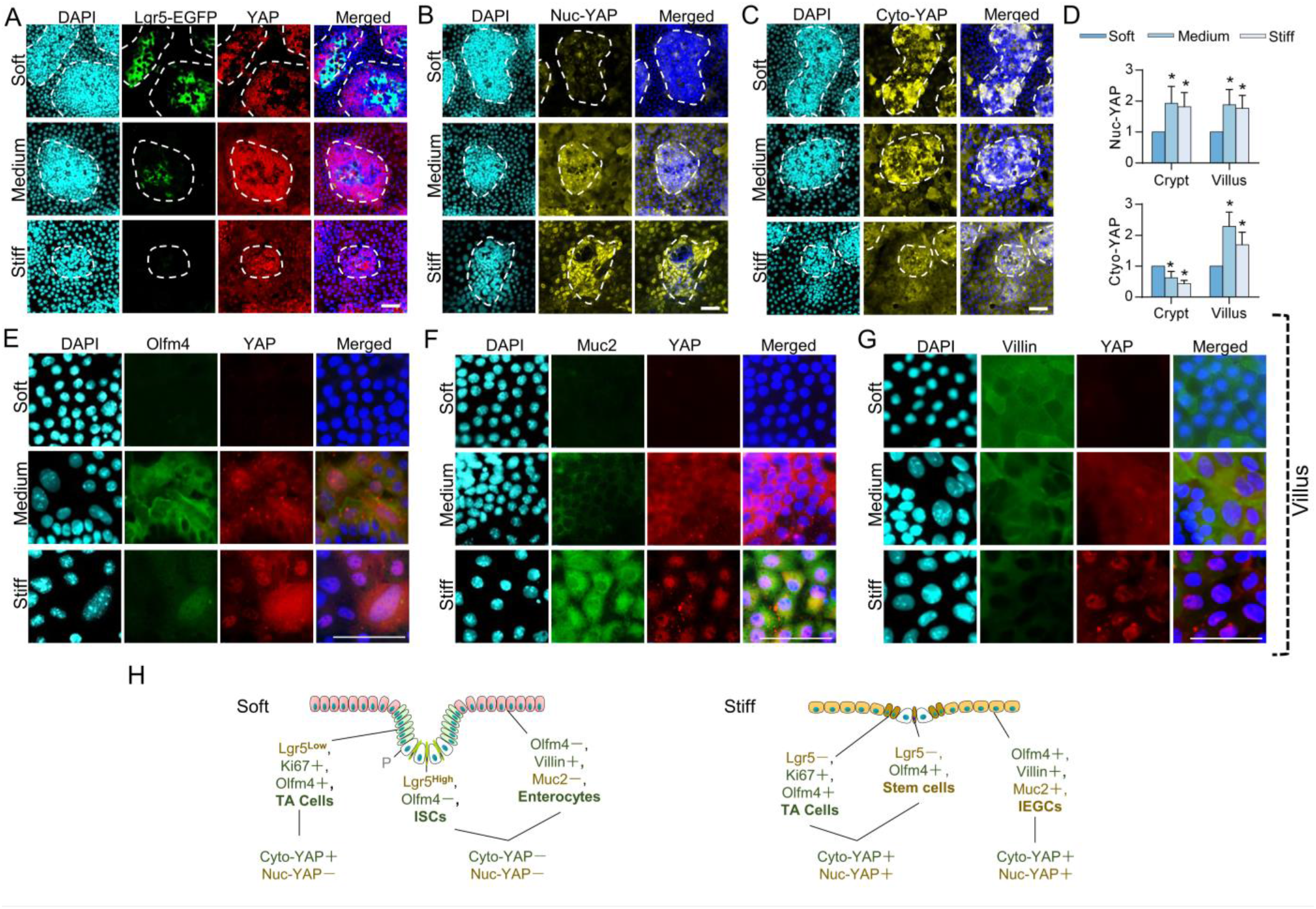
Stiffness regulates the fate of ISCs via YAP. (A) Lgr5-EGFP^high^ ISCs were YAP^−^ and disappeared when YAP was nuclear co-localized on the stiff matrix. The white dashed lines trace the crypt-like regions (*n*=3). (B) The non-phosphorylated nuclear (nuc-) YAP was increased by stiffening and showed clear nuclear co-localization on the stiff matrix (*n*=5). (C) The Ser 127 phosphorylated cytoplasmic (cyto-) YAP was decreased by stiffening in the crypt-like regions, but increased in the villus-like regions, which were quantified via fluorescent intensity (D, * V.S. Soft, *P*<0.05, *n*=5). (E) Olfm4 was highly expressed together with cyto-YAP (*n*=3). (F) Muc2 was highly expressed in the YAP nuclear co-localized cells (*n*=3). (G) Villin was highly expressed in the YAP^−^ cells (*n*=3). (H) The patterns of YAP were mapped onto all the cell types that were negatively (Green) or positively (Yellow) correlated with YAP nuclear translocation. Scale bar, 25 μm.

Based on the expression patterns of the nuc- and cyto-YAP (Fig. 2) we hypothesized that they play divergent roles in regulating the ISC differentiation patterns, where nuc-YAP appears to drive the differentiation into goblet cells, while cyto-YAP drives the differentiation into TA cells, enterocytes, and Olfm4^+^ stem cells (Fig. 2H). More specifically, on the soft matrix when the YAP^−^ and Lgr5^high^ ISCs migrate up, high cyto-YAP expression drives their differentiation into Ki67^+^ TA cells (Extended Data Fig. 9). Continuously driven by cyto-YAP, the TA cells primarily mature into enterocytes, which lose YAP expression after maturation (Fig. 2G). On the stiff matrix the constitutive expression of YAP causes the loss of Lgr5^high^ ISCs (Fig. 2A) and gain of Olfm4 instead (i.e., nuc-YAP^+^, cyto-YAP^+^, and Olfm4^+^ stem cells). These Olfm4^+^ stem cells have the potential to simultaneously differentiate into enterocytes driven by cyto-YAP and into goblet cells driven by nuc-YAP, which results in the new cell type, IEGCs (Fig. 2H).

## Nuc-YAP and cyto-YAP play divergent roles in determining the fate of ISCs

To test our hypothesis, transgenic mouse models were employed to knockout (KO) or overexpress (OE) YAP. Verteporfin (VP) was used to suppress YAP nuclear translocation ^17^. YAP KO led to the loss of the villus-like regions (Fig. 3A), indicating the indispensability of YAP in the differentiation of ISCs and the generation of villi. The leftover crypt-like regions were enriched with Paneth cells and were negative for nuc-YAP and Muc2, as well as cyto-YAP, Olfm4, Villin, and Egfr (Extended Data Fig. 12). YAP OE induced by doxycycline (DOX) led to the disappearance of the large and dense crypt-like regions on the soft matrix (Fig. 3B), causing a shift towards the stiff matrix-like phenotypes. Conversely, VP administration on the stiff matrix led to the formation of large crypt-like regions (Fig. 3C) and the restoration of Lgr5 expression (Fig. 3D), bestowing soft matrix-like phenotypes.

**Figure 3.**
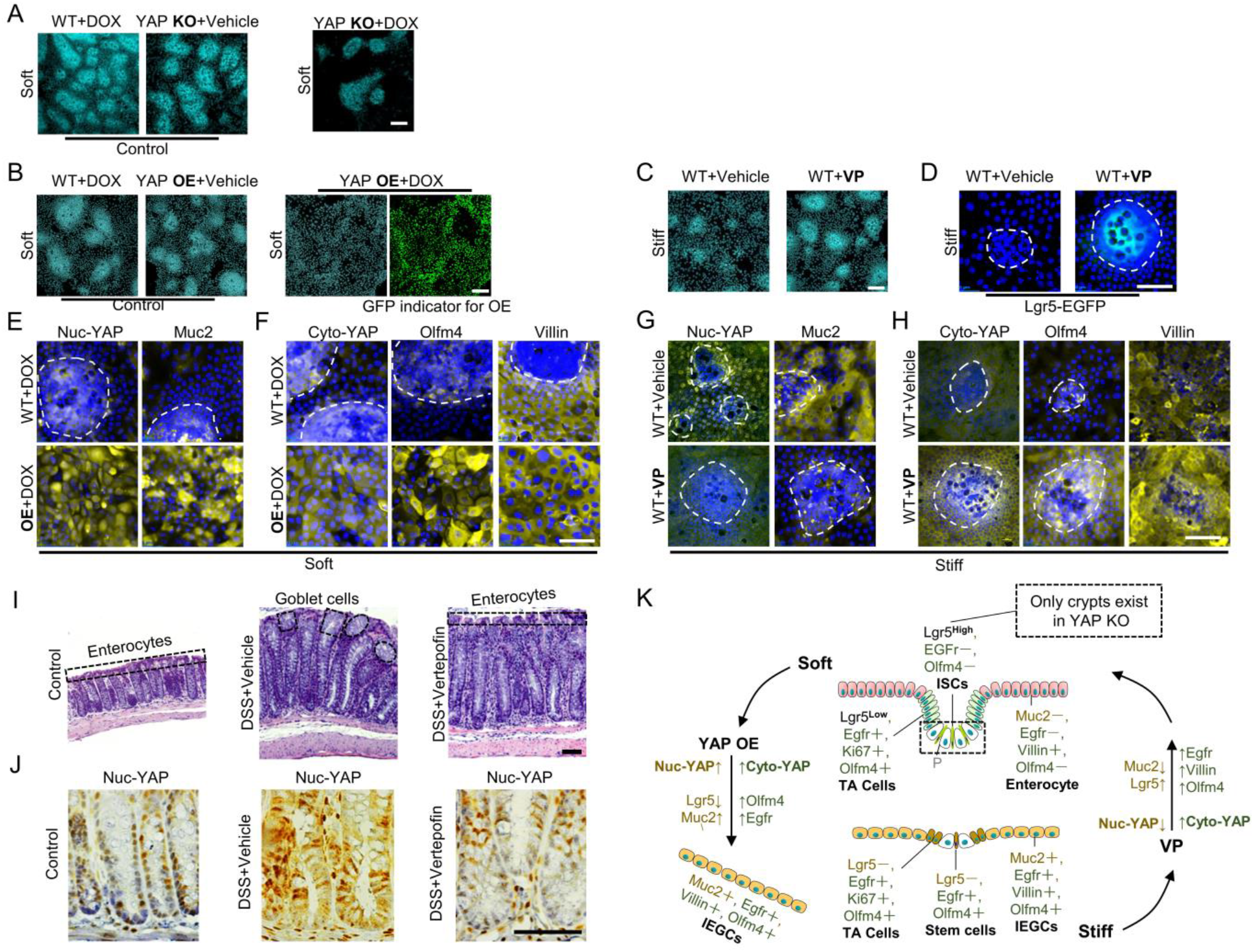
The fate of ISCs was manipulated via YAP knockout (KO), overexpression (OE), and Verteporfin (VP). (A) The villus-like regions vanished in the YAP KO groups. (B) YAP OE led to the loss of the crypt-like regions on the soft matrix. (C and D) VP administration increased the size of the crypt-like regions and resumed the Lgr5-EGFP expression on the stiff matrix. (E) Increase of nuc-YAP via OE augmented Muc2. (F) Increase of cyto-YAP via OE augmented Olfm4. No significant changes of Villin were detected. (G) Decrease of nuc-YAP via VP suppressed Muc2. (H) Increase of cyto-YAP via VP augmented Olfm4 and Villin. (I) Goblet cells replaced the enterocyte in the colon brush border in DDS-induced colitis group, and VP reversed this replacement. (J) Nuc-YAP was increased in the DDS-induced colitis group, but VP suppressed nuc-YAP. (K) YAP OE transformed the soft-matrix phenotypes into the stiff-matrix phenotypes, and VP did the opposite. ‘Yellow’ indicates the regulation by nuc-YAP. ‘Green’, the regulation by cyto-YAP. The white dashed lines trace the crypt-like regions. Scale bars in A, B and E, 100 μm; I and J, 200 μm; the rest, 25 μm. *n*=3 for these experiments.

Comparing the phenotypes between YAP OE and VP administration showed that nuc-YAP was increased in YAP OE but decreased in VP (Fig. 3E and 3G). In contrast, cyto-YAP was consistently increased in both models (Fig. 3F and 3H). Therefore, comparison of these two models enables us to discriminate the functional roles of nuc-YAP and cyto-YAP. Increasing nuc-YAP expression by OE promoted Muc2 (Fig. 3E), whereas decreasing nuc-YAP by VP suppressed Muc2 (Fig. 3G). Meanwhile, the increase in cyto-YAP by both OE and VP persistently augmented the expression of Olfm4, Villin, and Egfr (Fig. 3F and 3H, OE + Vehicle and Egfr in Extended Data Figs. 13 and 14). Nevertheless, YAP OE did not significantly increase Villin expression, which is most-likely because the basal level of Villin expression was saturated on the soft matrix (i.e., WT+DOX in Fig. 3D).

To assess the impact of tissue stiffness and VP *in vivo*, we administered VP in the dextran sulfate sodium (DSS)-induced chronic colitis mouse model (Method). In the colitis mouse the colon thickened (Fig. 3I and Extended Data Fig. 15A) and stiffened (Extended Data Fig. 15B). Moreover, colitis induced the replacement of enterocytes by goblet cells in the colon brush border, which was reversed by VP administration (Fig. 3I). Mechanistically, similar to the *in vitro* VP administration, the stiffened colon in the colitis mouse increased nuc-YAP which promoted the differentiation towards to goblet cells, and VP suppressed the stiffening-induced increase of nuc-YAP as well as goblet cell differentiation (Fig. 3I and 3J). The increase of cyto-YAP in the VP treatment group augmented the expression of Olfm4 and Egfr, which is also consistent with the *in vitro* observation (Extended Data Fig. 15D). Therefore, we demonstrated that YAP is indispensable for the ISC differentiation whereby nuc-YAP drives the differentiation towards Muc2^+^ goblet cells, and cyto-YAP drives the differentiation towards Villin^+^ enterocytes meanwhile promoting the expression of Egfr and Olfm4 (Fig. 3K).

### Stiffness-regulated transcriptional signatures

We generated, for the first time, the stiffness-regulated transcriptional signatures of the ISCs and their differentiated cells using single-cell RNA sequencing (scRNAseq) analysis of our quasi-3D organoids. The single-cell expression profiles were clustered into thirteen cell groups (Fig. 4A) and the most highly expressed genes of each group were easily distinguishable (Supplementary Table 1). Using the known marker genes ^18^, they were identified as various types of gut epithelial cells, matching the *in vivo* gut epithelium (Figs. 4A and 4B, and Extended Data Fig. 16). The clustering was consistent across both the soft and stiff matrix (Extended Data Fig. 17). Notably, two cell groups were identified as IEGCs which expressed mild levels of marker genes for stemness, enterocytes, goblet cells as well as other secretory cell types (Fig. 4B).

**Figure 4.**
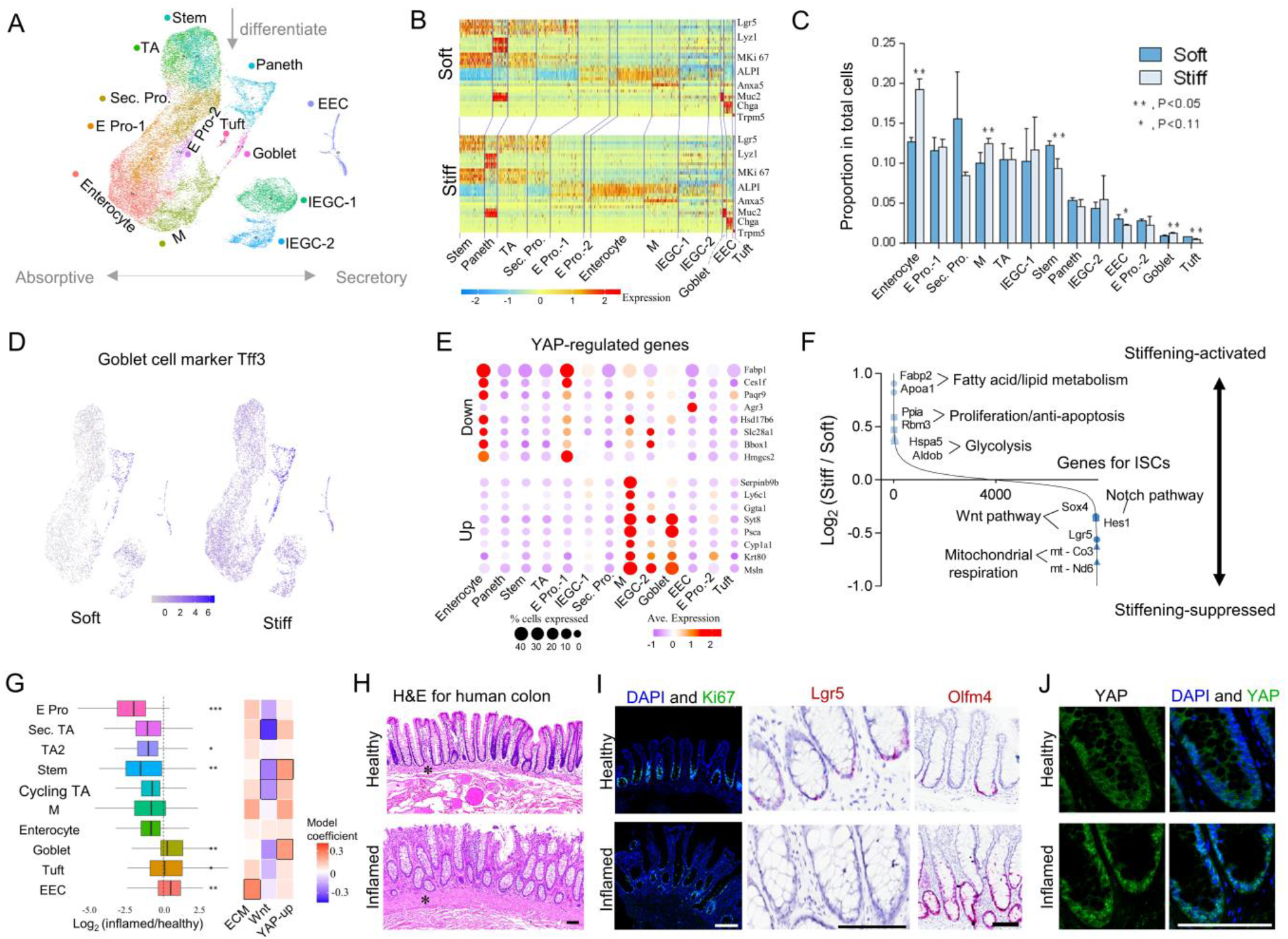
Single cell RNA sequencing and histology from IBD patient. (A) UMAP plot with the cell clusters (marked by color) including ISCs and the differentiated cells. ‘Sec’, secretory; ‘Pro’, progenitor; ‘E’, enterocyte; ‘M’, microfold. (B) Heat map for marker genes of each cell type (Extended Data Fig. 16 for full version). (C) The proportions of each cell type on the soft and stiff matrix. (D) Expression of Tff3 was higher on the stiff matrix than on the soft matrix. (E) The genes downregulated by YAP were highly expressed in enterocytes and E pro.- 1; however, the genes upregulated by YAP were highly expressed in goblet cells, IEGCs-1 and M cells. (F) Differential gene expression analysis in ISCs showed that stiffening suppressed both Wnt signaling (e.g., Lgr5 and Sox4 genes), and Notch signaling (e.g., Hes1), and possibly switched metabolic phenotype from mitochondrial respiration (e.g., downregulated mt-Co3 and me-No6) to glycolysis (e.g., upregulated Hspa5 and Aldob). *n*=3 for A-F. (G) Compared to healthy individuals (*n*=5), relative proportions of cell types in IBD patients (*n*=3) showed a decrease of ISCs, an increase of goblet cells and a trend towards a decrease of enterocytes. \**P*<0.05, \*\**P*<0.01, \*\*\**P*<0.001. Pathway enrichment analysis shows that in IBD patients ECM secretion is activated in EECs, Wnt signaling is suppressed in ISCs and the YAP up-regulated genes are highly expressed in ISCs and goblet cells. Model coefficients are output of linear mixed model from gene signatures associated with the respective pathways. Black outline for each box represents *P*<0.05 for linear mixed model and *P*<0.05 for pathway enrichment (Method). (H) H&E staining shows the thickening of BM and lamina propria labelled with asterisks, and the disappearance of the enterocyte brush border in the human inflamed colon. (I) Ki67^+^ proliferating cells and Lgr5^+^ ISCs were decreased, and Olfm4^+^ cells were increased in the inflamed colon. Lgr5 and Olfm4 were stained via in situ hybridization. (J) YAP showed more nuclear localization in the inflamed colon. *n*=3 for human colon resection samples. Scale bar, 200 µm.

The relative number of cells in each subset further confirmed the reduction in both ISCs and EECs, and the increase in goblet cells on the stiff matrix (Fig. 4C). Although stiffening appeared to increase the number of enterocytes (Fig. 4C), differential expression analysis (Supplementary Table 2) showed that the enterocytes on the stiff matrix expressed high levels of the goblet cell marker, Trefoil factor 3 (Tff3). In fact, Tff3 was upregulated across most of the cells on the stiff matrix (Fig. 4D), which was also confirmed at the protein level (Extended Data Fig. 18). These results suggest a preferential differentiation of ISCs towards goblet cells on the stiff matrix. In addition, the stiffening significantly increased M cells and decreased tuft cells (Fig. 4C).

We next assessed the expression of the YAP-up or down regulated genes (curated by Gregorieff et al. ^19^) in all the cell types. The YAP-downregulated genes (e.g., Fabp1 and Ces1f) were highly expressed in enterocytes and their progenitors, whereas the YAP-upregulated genes (e.g., Msln and Syt8) were enriched in goblet cells, IEGCs, and M cells (Fig. 4E and more YAP-regulated genes in Extended Data Figs. 19 and 20). These results further suggested that YAP could positively drive the differentiation of ISCs towards goblet cells as well as M-cells instead of enterocytes. Furthermore, the expression of the downstream genes of nuc-YAP, Id2, Birc5, and Areg ^20^ significantly increased on the stiff matrix (Extended Data Fig. 21). These results together with our experimental observations (Fig. 2F) indicate that nuc-YAP drives the ISC differentiation towards goblet cells.

Stiffening suppressed both Wnt signaling (e.g., Lgr5 and Sox4 genes) and Notch signaling (e.g., Hes1) in ISCs (Fig. 4F). Wnt is indispensable for maintaining the stemness of Lgr5^+^ ISCs ^21^. Therefore, the stiffening-induced suppression of Wnt signaling could mediate the loss of Lgr5^high^ ISCs. Furthermore, the suppression of the Notch pathway could drive the differentiation of ISCs towards goblet cells instead of enterocytes ^22^, which is also in accordance with the stiffening-enhanced goblet cell differentiation. In addition, the differential expression analysis suggests a stiffening-induced metabolic reprogramming of ISCs, including an increase in glycolysis and a decrease in mitochondrial respiration (Fig. 4F). Furthermore, compared to the soft matrix, pathways involved in carbon metabolism are more enriched on the stiff matrix (Pathway enrichment analysis in Extended Data Fig. 22). The mechanotransduction signaling pathways including actin cytoskeleton, focal adhesion and tight junctions were also upregulated on the stiff matrix, which could potentially contribute to YAP activation (Extended Data Fig. 22).

To determine the extent to which the stiffening-induced remodeling of the murine gut epithelium resembles that in human IBD patients, we analyzed the scRNAseq data of colon epithelium (generated by Smillie et al. ^3^) and colon resection samples, from IBD patients and healthy individuals. The human single cell profiles were clustered into ten epithelial cell subsets which were annotated using the known marker genes (tSNE in Extended Data Fig. 23A and marker genes in 23B). The relative proportions of each cell type demonstrate that UC is associated with a decrease in ISCs, an increase in goblet cells, and a trend towards a decrease in enterocytes (Fig. 4G and Extended Data Fig. 23C, with and without accounting for sample size, respectively). Pathway enrichment analysis demonstrates a strong activation of pathways involved in extracellular matrix biosynthesis (including collagen I and IV) in UC, indicating fibrosis and stiffening (Fig. 4G). Concomitantly, Wnt signaling was suppressed in the ISCs of UC patients (Fig. 4G and Extended Data Fig. 23D). The cluster of genes that have been previously shown to be directly upregulated by YAP ^19^ were highly enriched in both ISCs and goblet cells of UC-derived tissues, indicating the upregulation of YAP in these cells (Fig. 4G). In addition, there is a strong activation of the mechanosignaling pathway, including integrins, YAP, and TEADs (the primary transcription factors for YAP), in both the ISCs and the goblet cells of UC, but not in enterocytes (Extended Data Fig. 23E). Together, these results suggest a mechanosignaling-induced ISC reprograming in UC, where the BM stiffening-induced upregulated integrin signaling promotes the expression of YAP as well as its interaction with TEAD in the nuclei, which potentially suppresses Wnt signaling in ISCs and drives their differentiation towards goblet cells.

Human UC colon samples showed the disappearance of the enterocyte brush border and the thickening of BM and lamina propria (Fig. 4H). The excessive collagen deposition confirmed the occurrence of fibrosis (Extended Data Fig. 24A-C). Meanwhile, similar to our *in vitro* observations, Muc2^+^ goblet cells were overwhelmingly present in the inflamed colon compared to the healthy colon (Extended Data Fig. 24D). The number of Ki67^+^ proliferating cells and Lgr5^+^ ISCs were significantly decreased in the inflamed samples (Fig. 4I). Nevertheless, Olfm4 expression was increased and extended into larger regions (Fig. 4I and Extended Data Fig. 24E). Notably, there is a strong YAP nuclear localization in the inflamed samples which could be induced by BM fibrosis and stiffening (Fig. 4J). In the extremely fibrotic strictured ileum of CD patient, the samples exhibited a complete loss of the normal invaginated crypt structures (Extended Data Fig. 25). Meanwhile, a large number of ectopic crypts were formed in the hyperplastic BM and lamina propria. These phenotypes of the strictured ileum closely resemble the *in vitro* stiffening-induced decrease of crypt size and the formation of ectopic crypts (Fig. 1G). The scRNAseq and histology data from human IBD patients, which strongly complement our *in vitro* observations, demonstrate that gut fibrosis is associated with a strong YAP nuclear translocation, loss of ISCs, extension of Olfm4, and enhanced differentiation towards goblet cells, all of which we have shown to be a direct consequence of tissue stiffening.

## Discussion

We have developed a novel culture of quasi-3D gut organoids plated on top of soft hydrogel matrix. The resulting *in vitro* cellular collective spontaneously reorganizes into a structure reminiscent of the crypt-villus anatomy and diverse cell types of native *in vivo* gut epithelium. While the 3D gut organoid cultured in Matrigel^®^ has widely been used as an *in vitro* model of intestinal epithelium, it faces key limitations. Most notably, its intrinsic configuration does not mimic the *in vivo* anatomy and instead more closely resembles tumor conditions. Furthermore, the temporal and spatial nutrient and pressure gradients inside the Matrigel^®^, could directly impact the growth of 3D organoids which underlie a substantial experimental variability ^23^. By contrast, the novel system described here effectively circumvents these limitations and spontaneously generates crypt and villus structures with complex cellular composition similar to those *in vivo*.

Using this system, we observed that increasing matrix stiffness significantly reduced the TA progenitor cells and Lgr5^high^ ISCs, while driving their differentiation preferentially towards goblet cells. These results were confirmed in colitis mice and IBD patients. It is notable that it was previously reported that when the 3D gut organoids were embedded inside of the matrix, the matrix stiffening helped maintain the ISCs and suppressed their differentiation ^24^. This discrepancy might be explained by the fact that the 3D organoids more resemble the tumor microenvironment. Additionally, our data show that the increased expression of nuc-YAP on the stiff matrix caused the reduction of ISCs, which is in-line with the results using Lats1/2 knockout mice ^25^. We also elucidated, for the first time, the divergent roles of cyto-YAP and nuc-YAP in determining the direction of ISC differentiation. On the soft matrix, cyto-YAP dominates ISC differentiation into mature enterocytes, while on the stiff matrix, the nuc-YAP drives the differentiation towards goblet cells, giving rise to the new cell type, IEGC. Our data also have direct implications for the role of BM stiffening and YAP signaling in the progression of IBD to carcinoma. As suggested by our observations and the previous studies ^26, 27^, increasing YAP expression could decrease the incidence of Lgr5^+^ ISCs to form tumor. Nevertheless, the matrix stiffening expanded the expression of cyto-YAP from crypt-like regions into villus-like regions, which, in turn, expanded the expression of the oncogenes, Olfm4 and Egfr ^28–30^. Moreover, the stiffening led to the formation of new crypts in the villus-like regions, which could contribute to the uncontrolled regeneration of gut epithelium and the development of carcinoma.

## Acknowledgements

This work was supported by funding from National Institutes of Health (R01DK123219, R01EY028234, and K01DK103947 to N.S., and R01HL148152 and U01CA202123 to J.J.F.), ECOR/MGH (2019A002949 to N.S.), and Polsky family fund (to N.S.). We thank Dr. Ramnik J. Xavier, the Director of the Center for the Study of Inflammatory Bowel Disease at Massachusetts General Hospital (National Institutes of Health, P30DK043351) for his constructive comments, providing human samples and sharing the human scRNAseq data. We thank Maris A. Handley and Jacqueline Choi from the HSCI-CRM Flow Cytometry Core and iHisto Inc for their supports.

## Author Contributions

S.H. and N.S. conceptualized the work and designed the experiments. S.H. and P.L. performed the experiments with inputs from J.W., P.C., J.O.R. and F.D.C. on generating transgenic mice, C.Y.P. and J.J.F. for live cell imaging, M.M. and Ö.H.Y. for flow cytometry, E.S. and D.T.B for training, and S.I. for in situ hybridization. S.H., H.W. and V.M. performed the DSS mouse model with inputs from Z.Z. for measuring thickness, and R.W. and Y.Z. for measuring stiffness. W.K., K.S., T.X. and P.B. performed the scRNAseq analysis for human and organoids with guidance from S.H., R.S. and N.S.. S.H. and N.S. wrote the manuscript. J.J.F., F.D.C., P.C., M.M.,C.Y.P., D.T.B., T.X., M.H. and I.A. commented on the manuscript.

## The authors declare no competing financial interests

N.S. and S.H. are inventors on a patent application filed based on this investigation.

**Correspondence** and requests for materials should be addressed to N.S.

**Supplementary Information** is available for this paper.

## Methods

Mice including the strains of wild type C57BL/6J, *Lgr5-EGFP-IRES-CreER* ^31^, YAP conditional knockout and YAP conditional overexpression were used for harvesting crypts. Polyacrylamide hydrogel matrix was fabricated as described before ^32^. The crypts collected from the mouse small intestines were counted and seeded on the hydrogel matrix. After 11 days culture the cells were fixed for immunofluorescent staining, or the cells were trpsinized for flow cytometry, the transferring to 3D organoid culture in Matrigel, and single cell RNA sequencing. The fluorescent signals after staining were quantified using a customized MATLAB code. Initial processing of scRNA-Seq sequencing data was performed using CellRanger (v4.0.0). Further analysis was performed using the Seurat and Phenograph. C57BL/6J mice were given 3 cycles of DSS to induce chronic colitis. The mouse colon tissues were collected for immunohistochemistry and the tissue stiffness was measured using an Instron uniaxial tester. The human samples were provided by the Prospective Registry in IBD Study at MGH (PRISM).

## Detailed Methods

### Mice

10-14 weeks old mice including the strains of wild type C57BL/6J, *Lgr5-EGFP-IRES-CreER* ^31^, YAP conditional knockout and YAP conditional overexpression were used for harvesting crypts. To generate the YAP conditional knockout mice, CAG-rtTA3 (Jackson Laboratories) mice were mated with (TetO)7-Cre (Jackson Laboratory) and Yap^fl/fl^ mice ^33^. To generate the YAP conditional overexpression mice, the tetO-YAP-GFP mice (Jackson Laboratory, 031279) expressing mutant S112A YAP crossbred with Villin-rtTA*M2 mice (Jackson Laboratory, 031285). 1 μg/ml DOX was added to induce YAP knockout or overexpression in the organoid culture.

### Preparation of the hydrogel matrix

Polyacrylamide (PA) hydrogel matrix were fabricated on a 35 mm dishes with glass bottom (Cellvis, D35-20-1.5-N) as described ^32^. Briefly, the recipe for different Young’s modulus were 3% acrylamide (Bio-Rad, 1610140) and 0.058% bisacrylamide (Bio-Rad, 1610142) for 0.6 kPa, 7.5% acrylamide and 0.034% bisacrylamide for 2.4 kPa, and 7.5% acrylamide and 0.148% bisacrylamide for 9.6 kPa. 0.1% ammonia persulfate (sigma, 09913) and 0.05% TEMED (Bio-Rad, 1610800) were added to start the polymerization process. Then, 70 μl gel solution was added into the dishes, and a cover slip of 18 mm in diameter was placed on the gel solution to flatten the surface and resulted in a gel of 300 μm in thickness (VWR, 48380 046). The polymerization required 40 minutes to 1 hour. Then, sulfo-SANPAH (Proteochem, C1111) was used to activate the gel surface under a 15 W 365 nm UV (VWR, 95-0042-07/36575-050) for 10 minutes. After the activation, 200 μl 0.1 mg/ml type I collagen (Advanced biomatrix, 5022) was added onto the gel overnight to covalently attach to the gel surface for cell culture.

### Harvest of crypts

The proximal 12∼15 cm small intestines were collected from mice. The intestinal lumen was opened longitudinally. The mucous was removed using the back of blades. Then, the intestine was washed with ice cold PBS without calcium and magnesium (Corning, 21-040), and cut into 5 mm ∼ 1 cm fragments and placed into 50 ml conical tubes that were filled with ice-cold 50 ml of PBS/EDTA (10 mM, Thermo fisher, 15575020). The fragments were incubated and shaken on ice for 40 minutes, and washed with ice-cold 50 ml of PBS. Then, the fragments were vigorously shaken in 25 ml PBS and filtered twice through a 70 μm mesh (BD Falcon) into a 50 ml conical tube to remove the villi and tissue pieces. The crypts were mainly in the suspension which were centrifuged for 5 minutes at 100g. The crypt pellet collected here was then ready for seeding on the hydrogel.

### Culture of Crypts

The crypt pellets were suspended in the seeding media and counted using cytometer to control the crypt density as 10,000 /ml. 150 μl to 200 µl crypt suspension was added on to the PA gel in the 35 mm dishes. After 4 hours, the floating cells/clusters were washed with PBS (Corning, 21-040-cv). 1.5 ml ENR media/dish was added and changed every other day. To make the ENR media, advanced DMEM/F12 (Gibco, 12634-028) was supplemented with 50 ng/ml EGF (Peprotech, 315-09), 100 ng/ml Noggin (Peprotech, 250-38), 10% R-spondin conditional media (iLab in Harvard digestive center), 1% Glutamax (Gibco, 35050-061), 1% HEPES (Gibco, 15630-080), 0.2% Primocin (Invivogen, ant-pm-2), 0.2% Normocin (Invivogen, ant-nr-2), 1% B27 (Gibco, 12587010), 0.5% N2 (Gibco, 17502-048), and 1.25 mM N-Acetyl-Cystein (Sigma, A8199). To make the seeding media, the ENR media was supplemented with 3 μM Chir-99021 (Selleckchem, S1263) and 10 μM Y-27632 (Sigma, Y0503). We captured the phase images and the GFP images for Lgr5-EGFP fluorescent signal every 20 mins for days using 20X objective of Leica microscope (Leica DMI8) with a live cell imaging system. Depending on different purposes, 1 μg/ml DOX and 1 μM VP was, respectively, added to induce YAP KO or OE, or inhibit YAP nuclear translocation during the culture. After 10-11 days culture, cells were fixed for immunofluorescent staining, or collected using TrypLE Express (Invitrogen, 12-605-010) for different purposes including flow cytometry, the transferring to 3D organoid culture in Matrigel, and single cell RNA sequencing.

### Immunofluorescence (IF), immunohistochemistry (IHC) and In situ hybridization (ISH)

For *in vitro* IF, cells were fixed in 4% paraformaldehyde/PBS for 10 minutes and cold 70% ethanol for 30 minutes, blocked in 2% BSA for 30 minutes and permeabilized in 0.3% Triton X-100/PBS for 20 minutes. The cell layers were stained with primary antibody, then stained with secondary antibodies. Laser scanning microscopy images were captured by using the inverted confocal microscope (Nikon C2, 20X or 60X objective). The average intensity of the fluorescent signals in these images were then quantified using a customized MATLAB code which can identify the crypt-like regions based on the nuclei density (Extended Data Fig. 26). All the fluorescent images represented at least nine field views from three different animal (3 filed views/animal). Student’s *t*-test was used to determine statistical significance with a cut-off of *P*<0.05. *In vivo* IHC and IF was performed by iHisto Inc. Samples were processed, embedded in paraffin, and sectioned at 4 μm. Paraffin sections were then deparaffinized and hydrated. Antigen retrieval was achieved by boiling the sections in 10 mM sodium citrate. Sections were then washed with PBS three times, treated with 3% H2O2 for 15 min and 5% bovine serum albumin for 20 minutes. The sections were incubated with primary antibodies overnight at 4 °C. Subsequently, the sections were immunohistochemically stained with secondary antibodies for 50 min at room temperature. Rabbit primary antibodies were used for staining Villin (ab130751), Muc2 (ab76774, ab272692), Chromogranin-A (ab15160), Lysozyme (ab2408), non-phosphorylated YAP (ab205270), Trefoil Factor 3 (ab272927) and Egfr (ab40815), Ki-67 (cs9129), Olfm4 (ab105861, cs14369), Ser127 Phospho-YAP (cs13008), collagen I (ab34710) and collagen IV (ab6586). Mouse primary antibodies were used for total YAP (ab56701) and Villin (Proteintech, 66096-1-Ig). Goat anti rabbit/mouse Alexa Fluor 405, 488, 594 and 647 (Thermo fisher) were used as secondary antibodies. DAPI (Fisher scientific, D1306) and UEA-1 Fluorescein (Vector labs, FL-1061-5) were respectively used for staining nuclei and Paneth cells. Single-molecule in situ hybridization was performed using Advanced Cell Diagnostics (ACD) RNAscope 2.0 HD Detection Kit (Fast Red dye) for the probes of Lgr5 (ACD, 311021) and Olfm4 (ACD, 311041).

### Flow cytometry

The cells on the PA hydrogel were trypsinized using TrypLE for 10 minutes at 37°C. After collecting the cells, cold SMEM (1:5) was added to stop the trypsinization, and followed by centrifuging for 5 min at 300 g. The cell pellets were re-suspended and stained for 15 min on ice in 1ml antibody cocktail. To make the antibody cocktail, SMEM (Sigma, M8167) was supplemented by CD45-PE (1:500, eBioscience, 30-F11), CD31-PE (1:500, Biolegend, Mec13.3), Ter119-PE (1:500, Biolegend, Ter119), CD24-Pacific Blue (1:300, Biolegend, M1/69), EPCAM-APC (1:300, eBioscience, G8.8) and CD117(C-kit)-APC-CY7 (1:300, Thermo fisher, 47-1171-80). After the staining 10ml SMEM were added and the samples were centrifuged for 5 min at 300 g. The pellets were resuspended with 1 ml SMEM supplemented by 7-AAD (1:500, Thermo fisher A1310), and filtered through a 40μm mesh (BD Falcon) before cell sorting with a BD FACS Aria II cell sorter. ISCs were isolated as Lgr5-EGFP^high^Epcam^+^ CD24^low/−^ CD31^−^ Ter119^−^ CD45^−^ and 7-AAD^−^, TA progenitors were isolated as Lgr5-EGFP^low^ Epcam^+^ CD24 ^low/−^ CD31^−^ Ter119^−^ CD45^−^ and 7-AAD^−^, Paneth cells were isolated as CD24^high^ Sidescatter^high^ C-kit^+^ Lgr5-EGFP^−^ Epcam^+^ CD31^−^ Ter119^−^ CD45^−^ and 7-AAD^−^.

#### Culture of 3D organoids in Matrigel transferred from the 2D hydrogel

The cells on the hydrogel martix were trypsinized using TrypLE for 5 minutes at 37°C. The wells were sealed with lid, and the bottom of the wells were vigorously hit onto the table to detach the cells. After collecting the cells/clusters cold SMEM (1:5) was added to stop the trypsinization, and followed by centrifuging for 5 min at 300 g. The cell pellets were resuspended in ENR media and Matrigel (1:1, Corning 356231), then seeded 25 μl/well in a 48-well plates and put at 37°C for solidification. After 20 minutes, 150 μl/well ENR media was added, and changed every 3 days. The 3D organoids were imaged on day 5.

### Single cell RNA sequencing for the gut organoids

The scRNA-seq library construction was performed on the Chromium 10x instrument using Chromium single cell 3’ reagent v3.0 kits, followed by sequencing on Illumina HiSeq 2500 instruments, which resulted in approximately 160 million reads per sample. Initial processing of scRNA-Seq sequencing data was performed using CellRanger (v4.0.0) (https://support.10xgenomics.com/single-cell-gene-expression/software/overview/welcome). In brief, reads were aligned to mm10 mouse reference genome with the mapping rate of ∼70%, followed by the generation of read counts per gene in each cell. Further analysis was performed using the Seurat 3.2.3 package (https://satijalab.org/seurat/). We filtered out cells with <200 expressed genes and genes expressed in <3 cells, followed by the exclusion of cells with high content of mitochondrial transcripts (> 20% of total reads). Counts across all cells for each sample were normalized using NormalizeData function and the effect of the cell cycle was removed by regressing the difference out between the S phase and G2M phase from normalized data. Using the FindVariableFeatures function we selected 2000 features to be used in a Principal Component Analysis (PCA). UMAP dimensionality reduction and cell clustering were performed using RunUMAP and FindClusters functions, respectively. VlnPlot and FeaturePlot functions were used to generate violin plots and feature plots for the datasets. Heatmaps of gene expression were generated using DoHeatmap function. Cells from all samples were then integrated using Seurat canonical correlation analysis (CCA) method. Integration anchors were obtained using the FindIntegrationAnchors function and used to integrate individual datasets with IntegrateData function. Biological annotation of cell clusters was based on the expression of known cell type markers. For the differential expression analysis, FindMarkers function was applied to the integrated samples in order to identify differentially expressed genes between the cell subsets.

### Single cell RNA sequencing for human colon epithelium

#### Cell clustering and differential expression analysis

We re-analyzed scRNAseq data of colon biopsy specimen generated by Smilie et al ^3^ (raw data from https://portals.broadinstitute.org/single_cell/study/SCP259). From the dataset, we used epithelial samples containing healthy tissue and inflamed tissue samples from patients with ulcerative colitis. We followed the same procedure to identify cell clusters as outlined in Smilie et al using Seurat (https://satijalab.org/seurat/, v.3.2.3) and Phenograph. The only exception being a larger k=1000 was used when applying Phenograph to KNN-graphs, and then re-clustering with k=50 is used to identify rare epithelial cell types. Cell clusters were identified by gene expression with known cell type markers. Barnes-Hut t-Distributed Neighbor Embedding on PCS (perplexity=10, iterations=5000) provided visualization of data embedding. The coarser k resulted in larger cell clusters where immature forms of cell types are no longer differentiated from the terminally differentiated cell types. The MAST model is fit to identify cell type markers and DE genes in inflamed tissue samples with control tissue, discrete coefficient of MAST model output is reported in text and figures unless otherwise stated.

#### Identifying statistically significant differences in cell proportions

As samples with exceedingly small number of cells show few cell types and disproportionate cell type proportions, we excluded samples containing less than 250 cells from subsequent analysis. Then changes in cell proportion between healthy and inflamed tissue are assessed by two methods. We first used monte-carlo test, where H0 is differences in proportions for each cell type between inflamed and healthy condition is a consequence of random sampling. We combined cells from both conditions together, and then randomly segregated cells back into the two conditions while maintaining original sample sizes and repeated the process 1000 times. We recalculated the proportion difference between the two conditions and compare to observed proportional difference for each cell type, and the *P*-value reflect number of simulations where simulated proportional difference was more than observed. This test reflects how enriched each cell type is within each condition (healthy or inflamed), but does not account for the specimen from which each cell is isolated. In the second method, we calculated the relative variation in each cell type proportion between all pairs of healthy donors as a control. Then, we calculated the relative variation in each cell type by dividing the fraction of the cell type in each inflamed specimen by that of a healthy specimen. After log2 transformation, we conducted a two-sided Kolmogorov-Smirnov (KS) test of the relative variation in composition between the control (healthy) and inflamed groups (ks.test function).

#### Identifying significant changes in pathway gene signatures

The ECM pathway (YAP1ECM_AXIS) was obtained from WikiPathways and WNT signaling pathway is obtained from KEGG. The gene signature of top 200 YAP-upregulated genes were curated by Gregorieff et al ^19^. Pathway enrichment analysis (PEA) were performed using gene sets from these pathways with the fgsea package in R, and the shared genes between significant cell types are used as the gene signatures for the pathways of interest. Expression of each gene was then scaled by its root mean squared expression across all cells, and mean scaled expression for all signature genes in the pathway is calculated to give a signature score for each cell. Then, we used mixed linear models to identify changes in expression levels of gene signatures in the inflamed state. In the model, the fixed effect term is used to represent the condition of each cell (healthy or inflamed) and the random intercept that varies with each sample is used to account for the sample each cell was isolated from. ANOVA is used to estimate the fixed term *P*-value.

### Chronic Colitis mouse model induced by DSS and samples from IBD patients

To induce chronic colitis, Wild type 8-weeks old C57BL/6J mice were given 3 cycles of DSS. Each cycle included 2.5% DSS in drinking water for 7 days and distilled water for the following 14 days. After the third cycle of giving DSS verteporfin (VP, 25 mg/kg/d in DMSO; Sigma-Aldrich) or vehicle as control was administered via i.p. injection for 14 days. Then colon tissues were collected for measuring stiffness and immunohistochemistry (IHC). Antibodies were used same as in IF. ID of human patients for the healthy, inflamed, and strictured gut samples are, respectively, 109922, 117324, and 110201. Protocol No. for involving human subjects is 2004P001067 reviewed by IRB.

### Uniaxial tensile testing for colon stiffness measurement

The colons collected from the chronic colitis experiments were cut open along the longitudinal direction into flat rectangular patches using a surgical scalpel. Sandpaper tabs were glued to both ends of a sample to prevent slipping during testing. The effective length (*l*_0_, i.e., sandpaper-to-sandpaper distance) and width (*w*) of each sample were measured using a caliper. The section view of the colons supported with a 10ul tips was imaged, and the thickness (*t*) of the section was measured using Image-J. Samples were mounted on an Instron uniaxial tester (Instron, Norwood, MA) by clamping the sandpaper tabs with the grips attached to the tester (Extended Data Fig. 27). Samples were moisturized with PBS spray before a test started. Steady-state uniaxial tensile tests were performed by fixing one end of the sample and pulling away the other end with an extension rate of 0.02 mm/s. The pulling force (*F*) and the displacement (*d*) of the moving end of the sample were recorded at a frequency of 5 Hz. The end of the regime of elastic deformation was marked by a drop in the slope of the force-displacement curve. The stain (*ε*) of the sample was obtained as

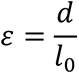

Assuming tissue incompressibility, the Cauchy stress (σ) can be calculated as

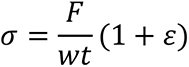

## Extended Data

**Extended Data Figure 1.**
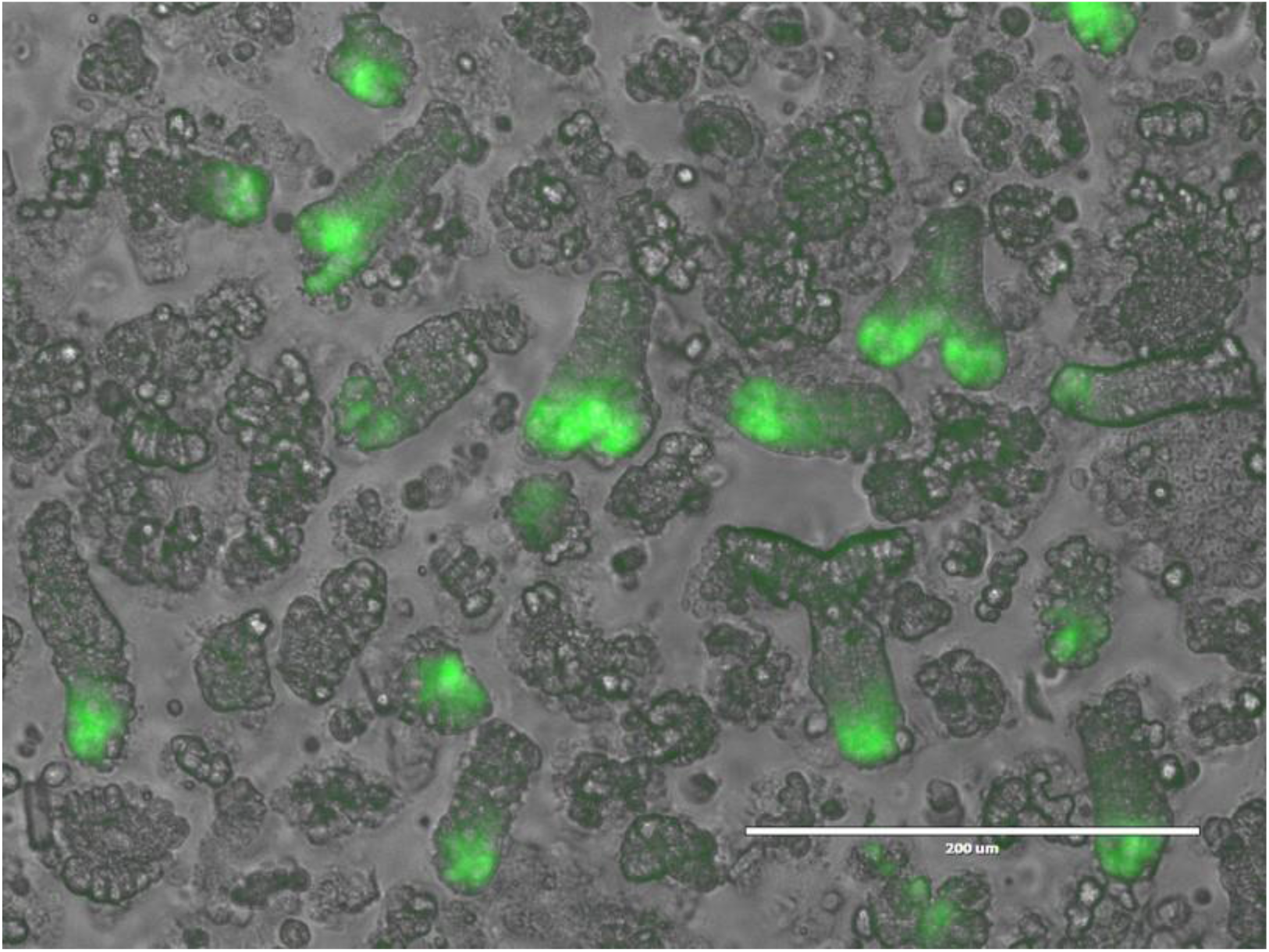
The crypts with Lgr5-EGFP^+^ ISCs were harvested from *Lgr5-EGFPIRES-creERT2* mice, and seeded on the hydrogel matrix.

**Extended Data Figure 2.**
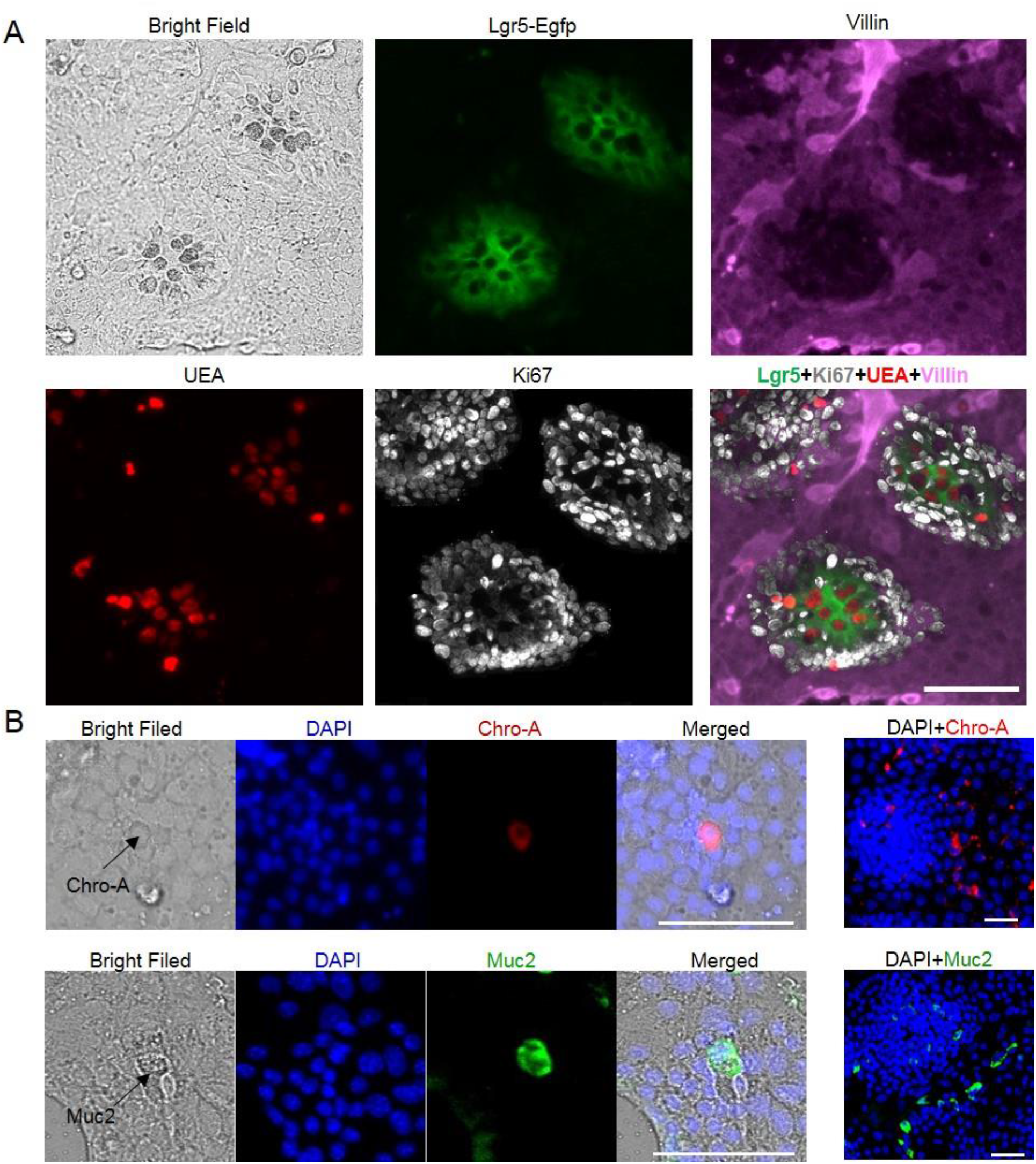
(A) Lgr5-EGFP^+^ ISCs were intermixed with the optically dark UEA^+^ Paneth cells, which were surrounded by Ki67+ TA cells in the crypt-like regions. Lgr5-EGFP^high^ ISCs weakly expressed Ki67. The villus-like regions were populated by Villin^+^ enterocytes. (B) Chro-A^+^ EEC and Muc2^+^ goblet cells were presented in the villus-like regions. *n*=3. Scale bar, 100 µm.

**Extended Data Figure 3.**
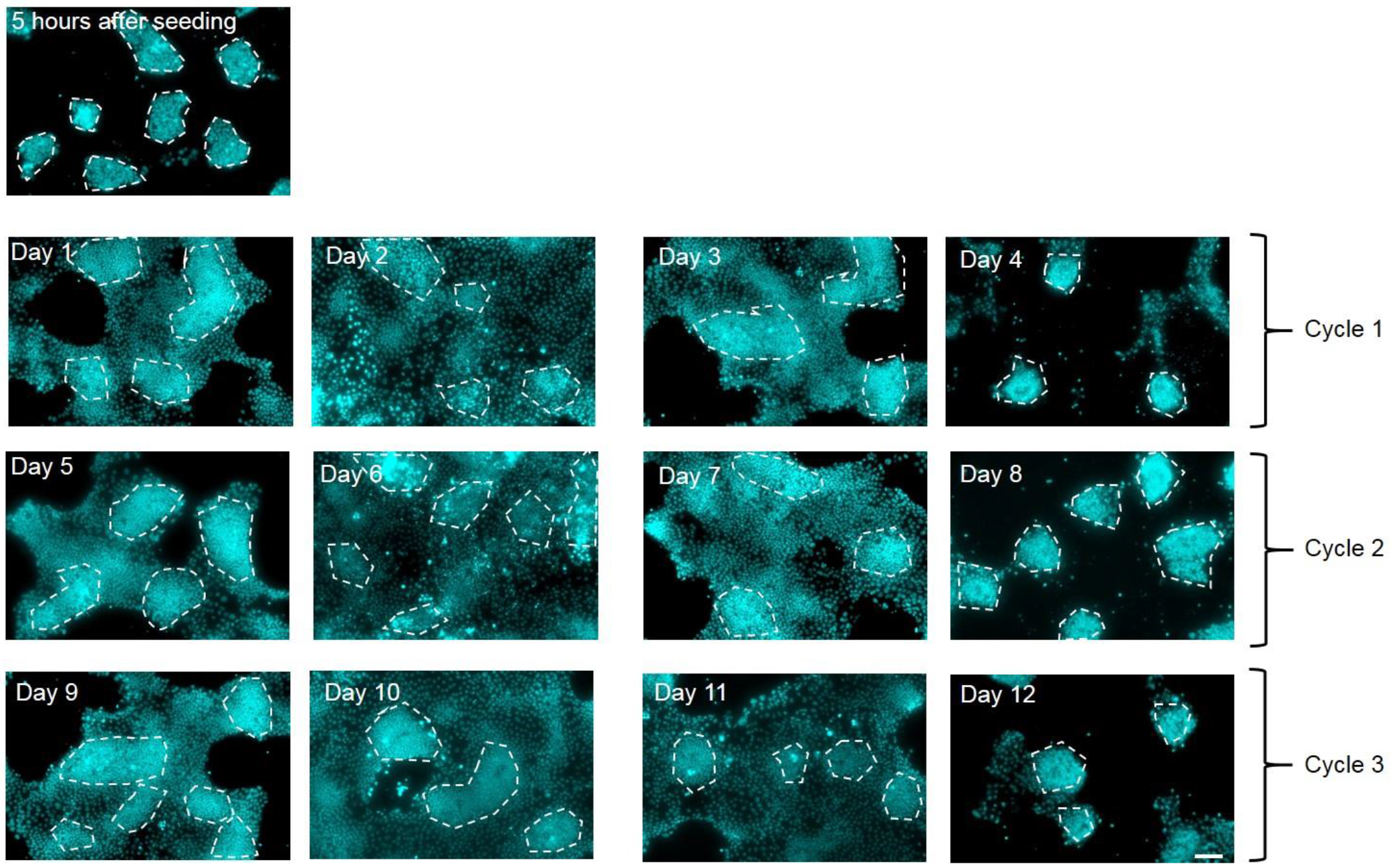
The villus-like regions exhibited a turnover rate of approximately 3 days (*n*=5). The white dashed lines trace the crypt-like regions. Scale bar, 100 μm.

**Extended Data Figure 4.**
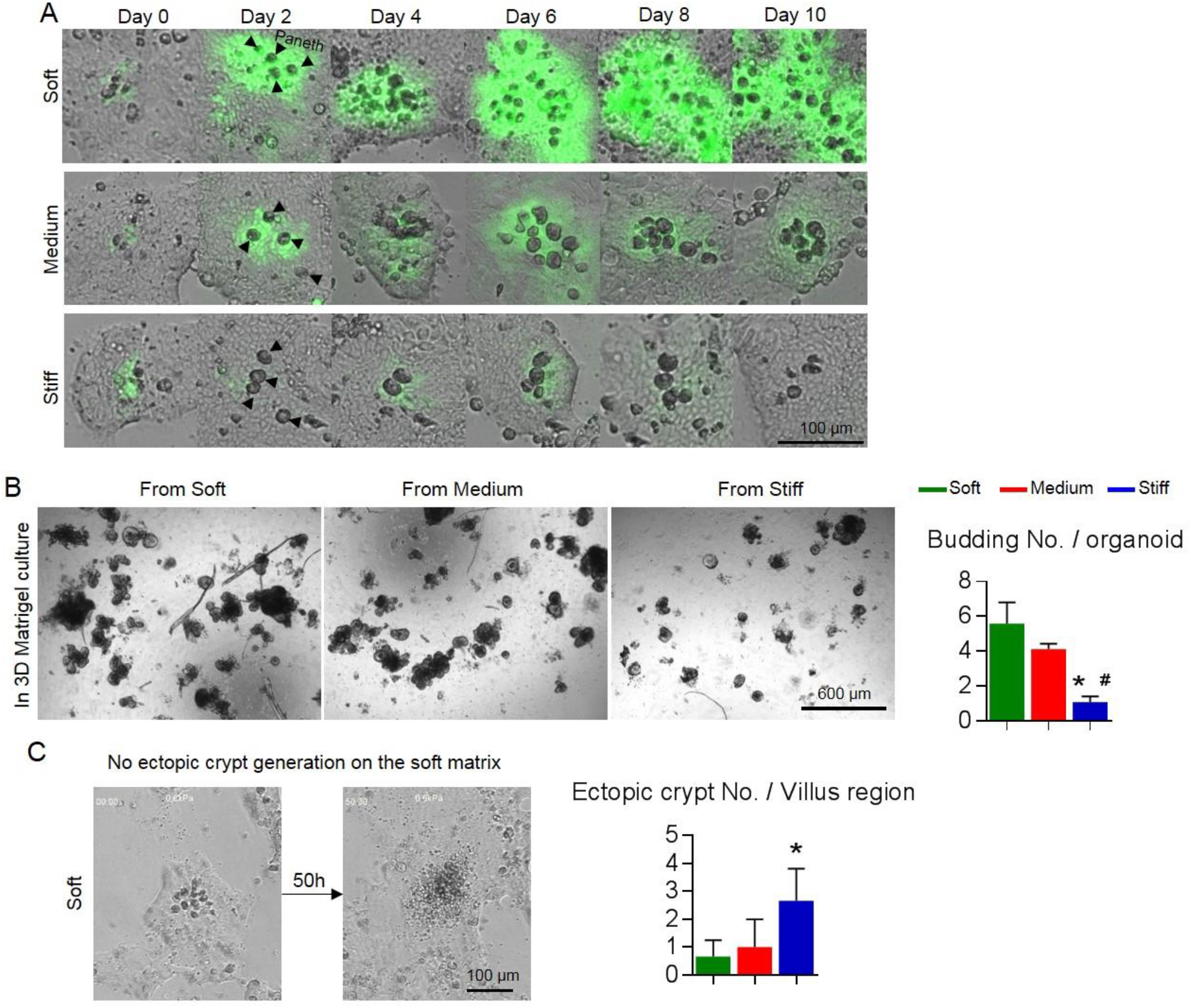
(A) Long-term live-cell imaging exhibited completely different phenotype of Lgr5-EGFP^+^ ISCs between the soft matrix and the stiff matrix. More specifically, on the soft matrix, the Lgr5-EGFP^+^ ISCs continuously increased and expanded throughout the culture period. In contrast, Lgr5-EGFP^+^ ISCs on the stiff matrix progressively diminished over time, nearly disappearing by the 10^th^ day. The medium matrix was able to maintain some Lgr5-EGFP^+^ ISCs. (B) The 3D organoids from the soft or medium matrix budded, but those from the stiff matrix grew as cysts with less budding (*n*=3). (C) New crypt generation in the villus-like regions appeared more on the stiff matrix than on the soft matrix (Movies. S2 and S3, *n*=3). * V.S. Soft and # V.S. Medium, *P*<0.05.

**Extended Data Figure 5.**
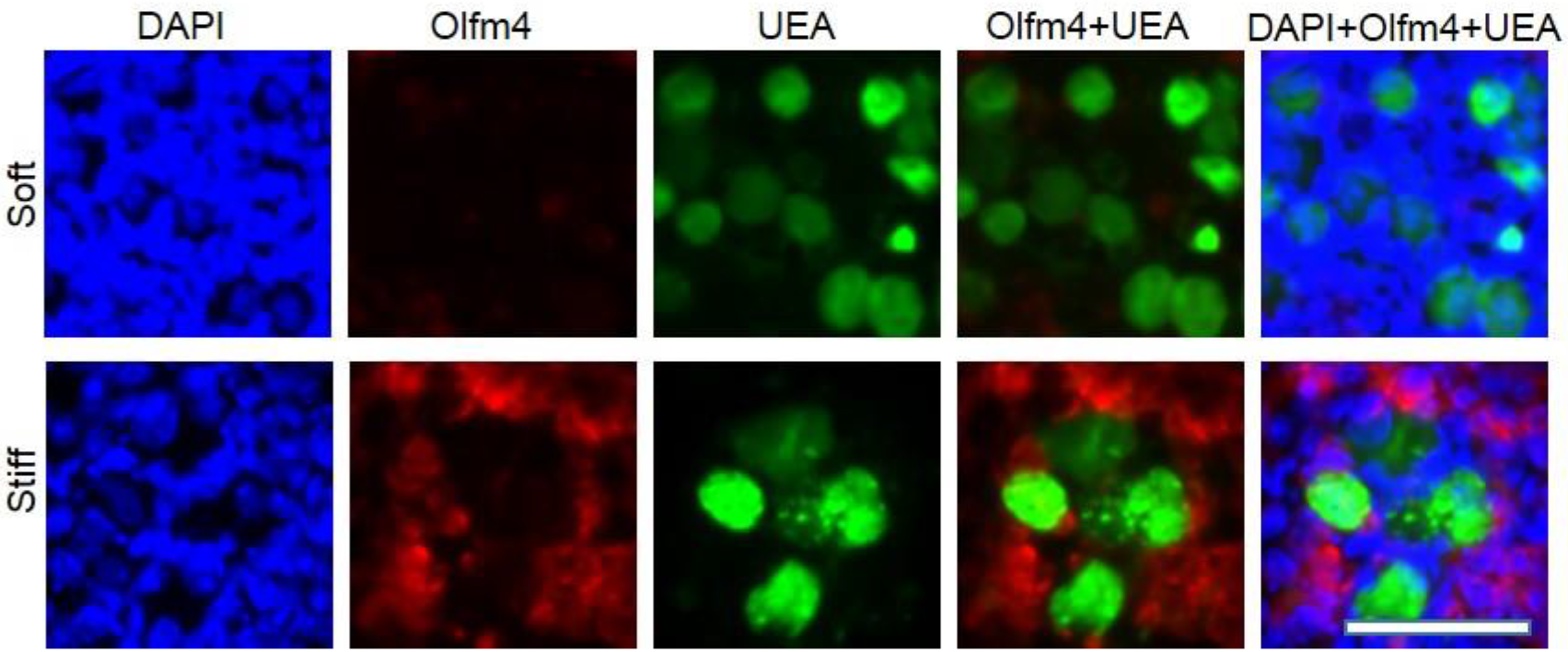
In the interior of the crypt-like regions, Olfm4^+^ cells became adjacent with UEA^+^ Paneth cells on the stiff matrix (*n*=3). Scale bar, 50 µm

**Extended Data Figure 6.**
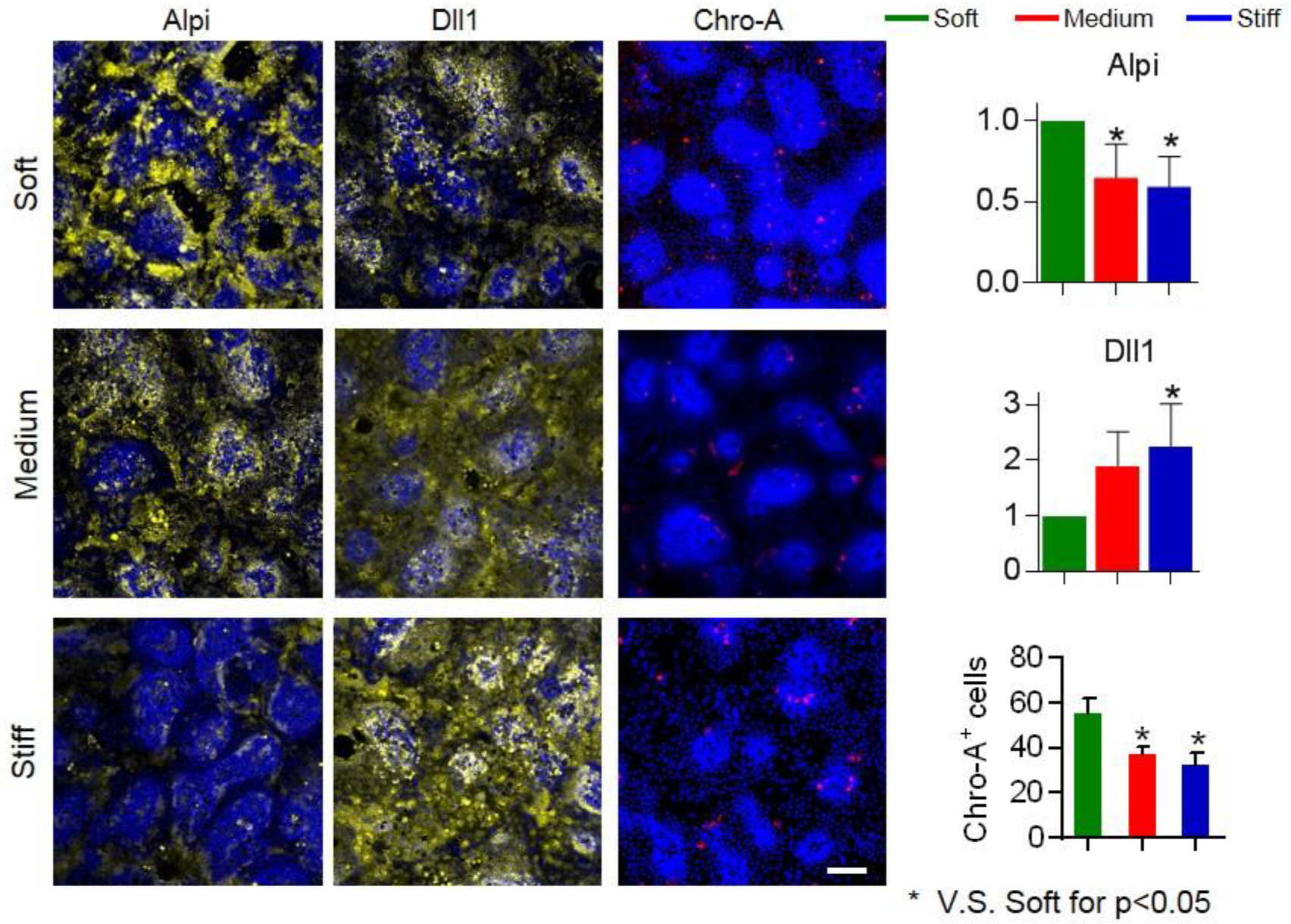
The stiffening decreased Alpi and Chro-A, and increased Dll1 (*n*=3). Scale bar, 100 μm.

**Extended Data Figure 7.**
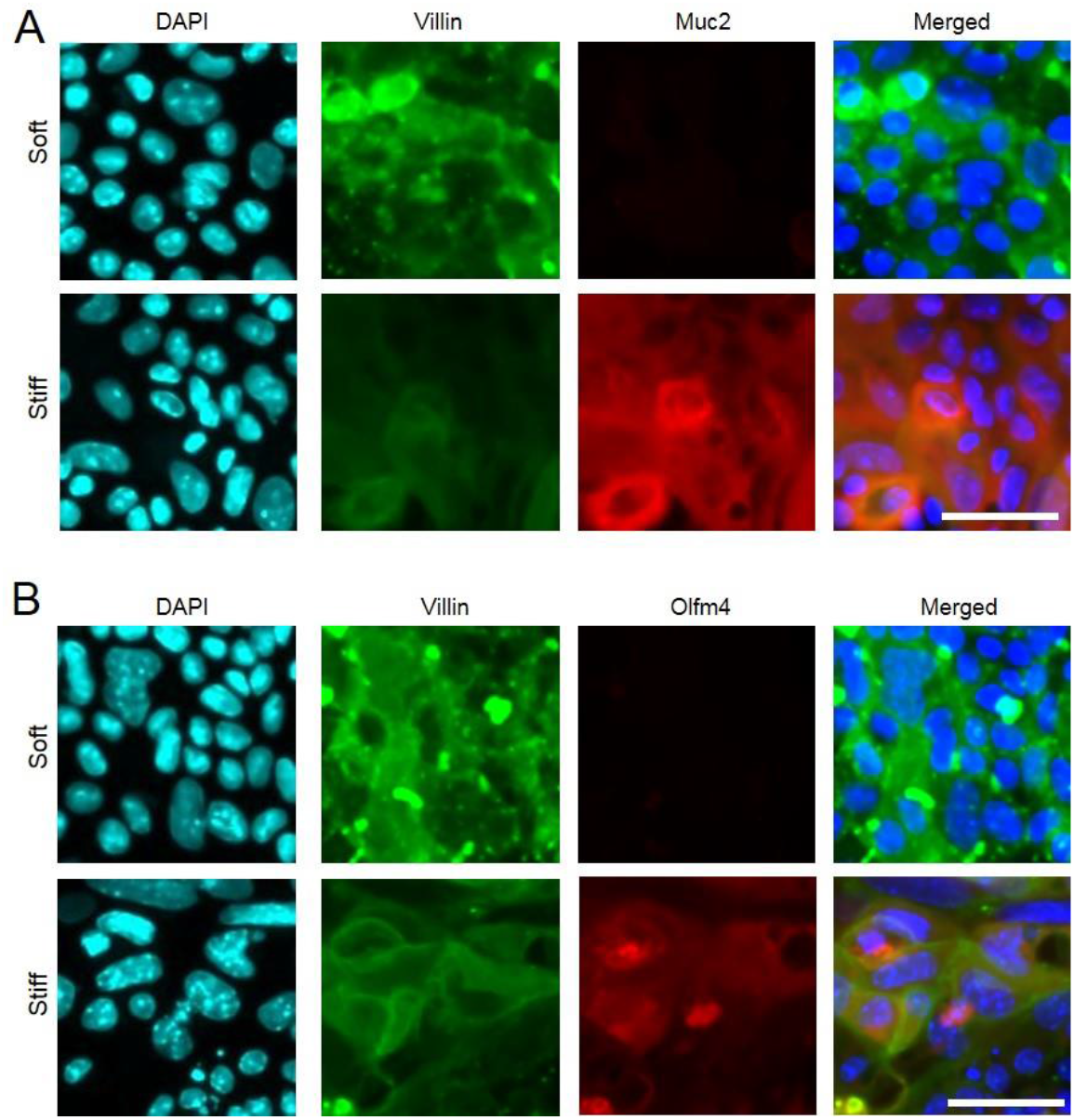
Counterstaining in the villus-like regions for Villin and Muc2 (A), and Villin and Olfm4 (B). *n*=3. Scale bar, 50 μm.

**Extended Data Figure 8.**
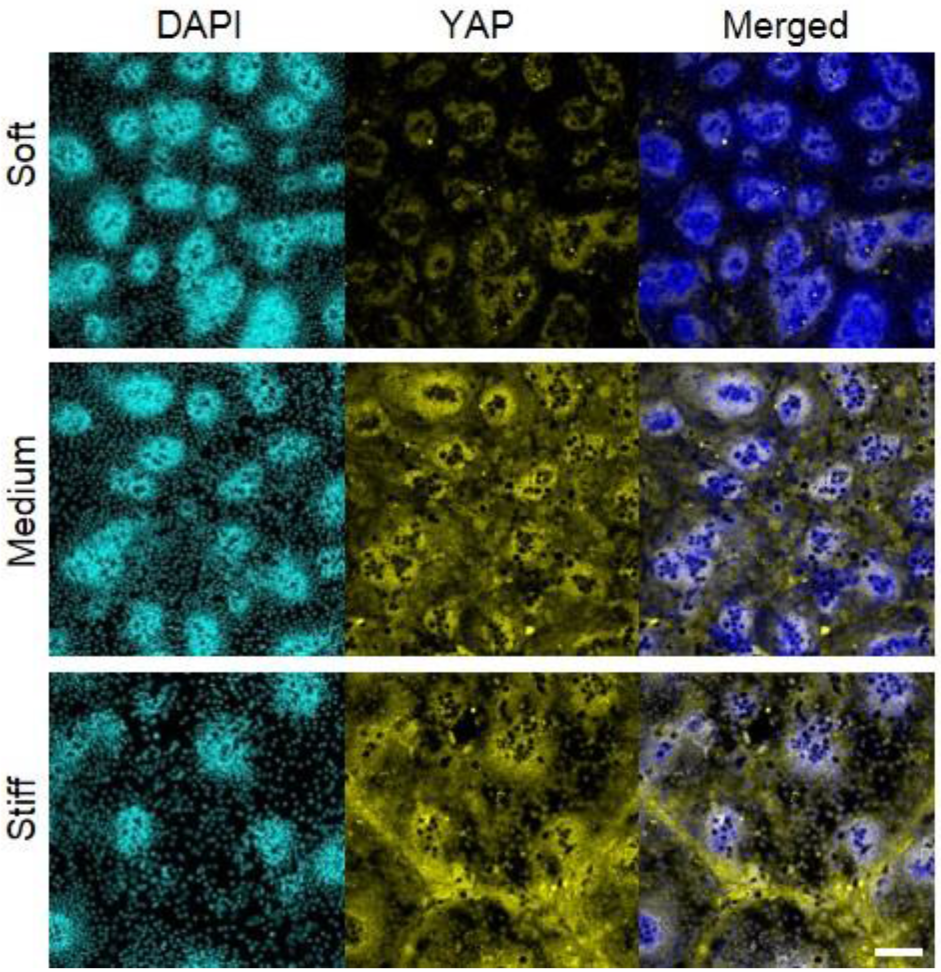
Stiffening increased YAP expression and promoted YAP nuclear translocation on the stiff matrix (*n*=5). Scale bar, 100 μm.

**Extended Data Figure 9.**
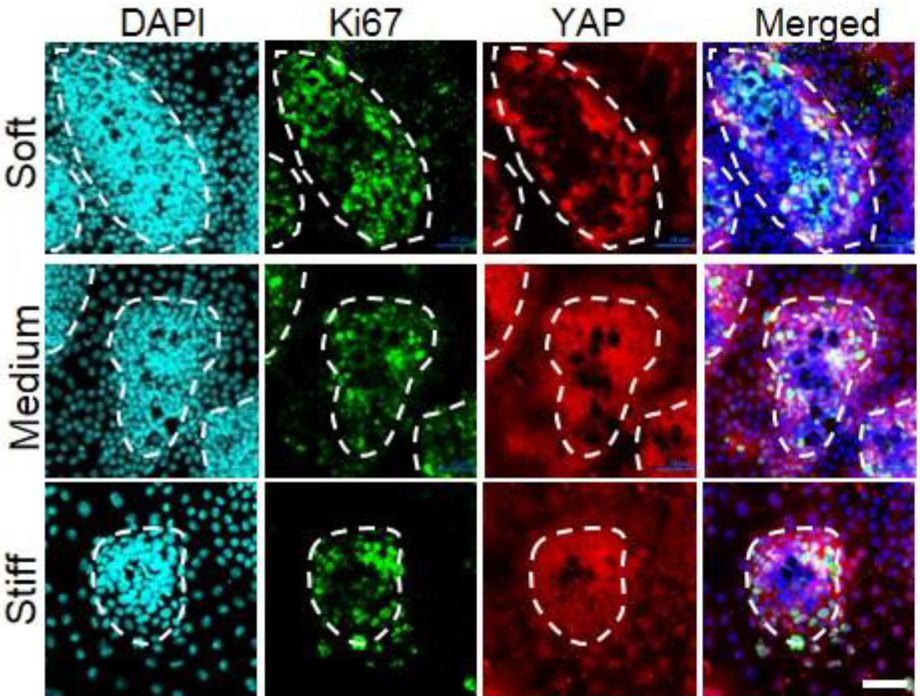
Ki67 was positively correlated with cyto-YAP (*n*=3). The white dashed lines trace the crypt-like regions. Scale bar, 25 μm.

**Extended Data Figure 10.**
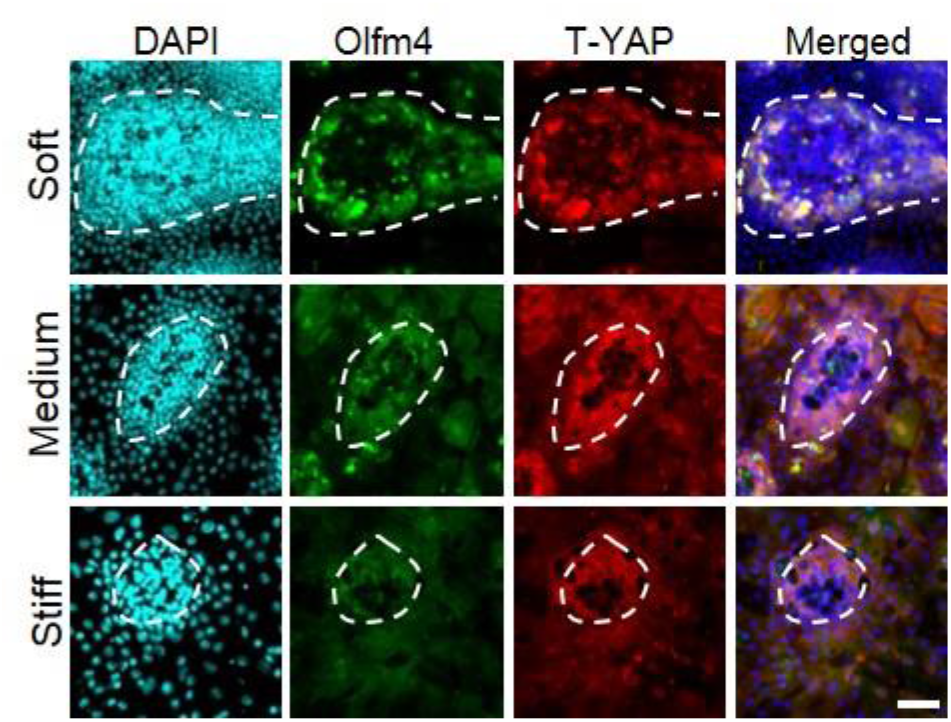
Olfm4 was positively correlated with cyto-YAP (*n*=3). The white dashed lines trace the crypt-like regions. Scale bar, 25 μm.

**Extended Data Figure 11.**
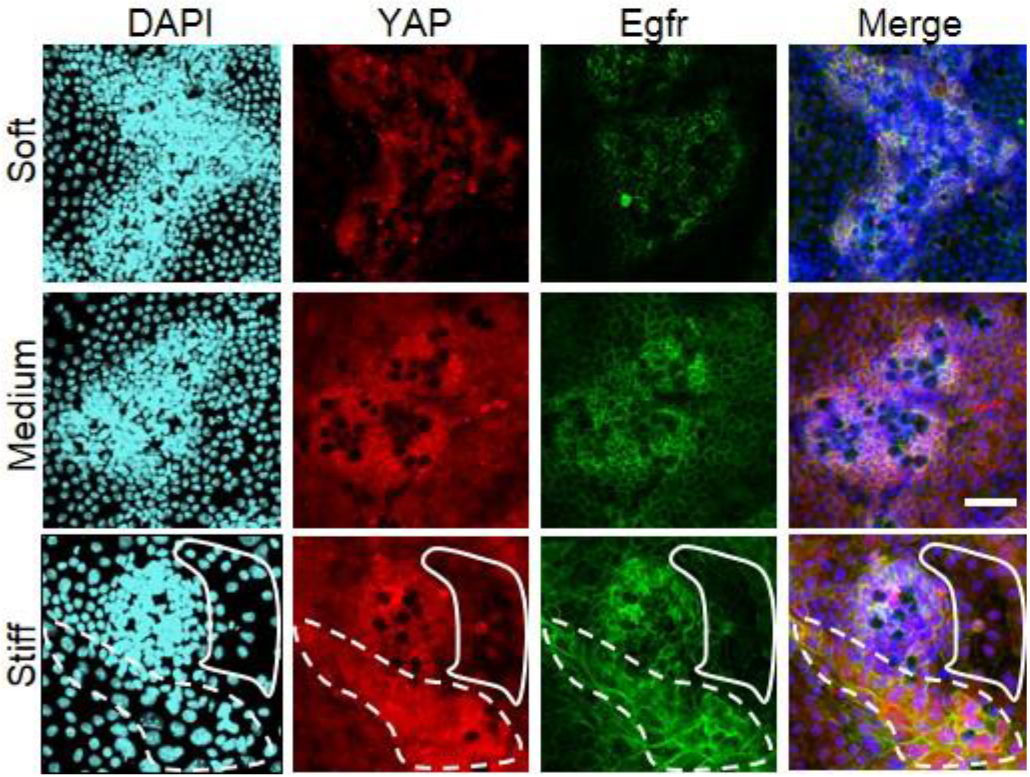
Egfr was positively correlated with cyto-YAP (*n*=3). On the stiff matrix, the white dashed line traces the region with high expression of cyto-YAP, and the solid line traces the region with YAP nuclear localization. Scale bar, 25 μm.

**Extended Data Figure 12.**
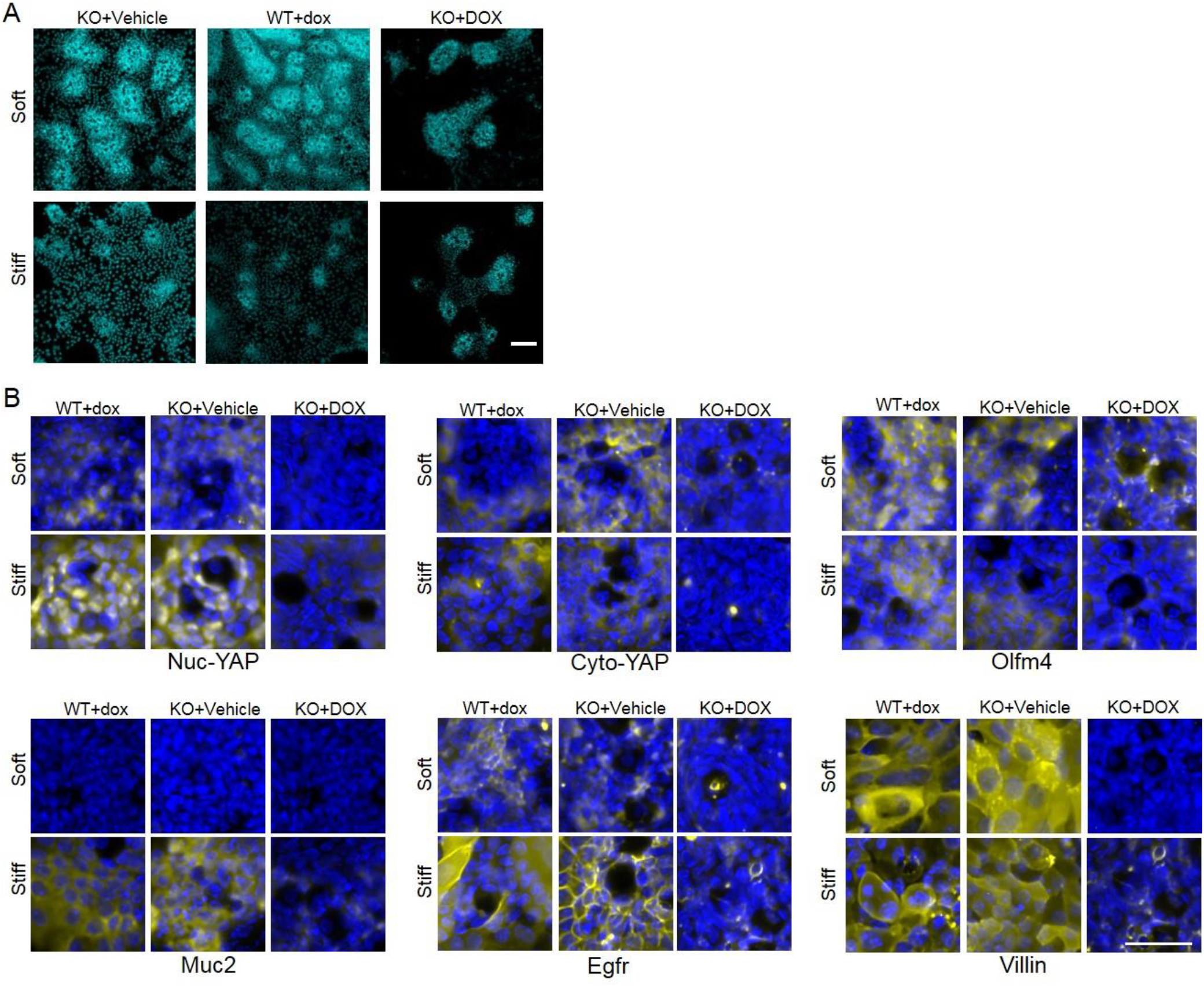
Staining for WT+DOX, YAP KO+Vehicle, and YAP KO+DOX on both soft and stiff matrix. (A) YAP KO by DOX led to the loss of the villus-like regions. Scale bar, 100 μm. (B) The leftover crypt-like regions were enriched with Paneth cells and were negative for nuc-YAP and Muc2, as well as cyto-YAP, Olfm4, Villin and Egfr. *n*=3. Scale bar, 25 μm.

**Extended Data Figure 13.**
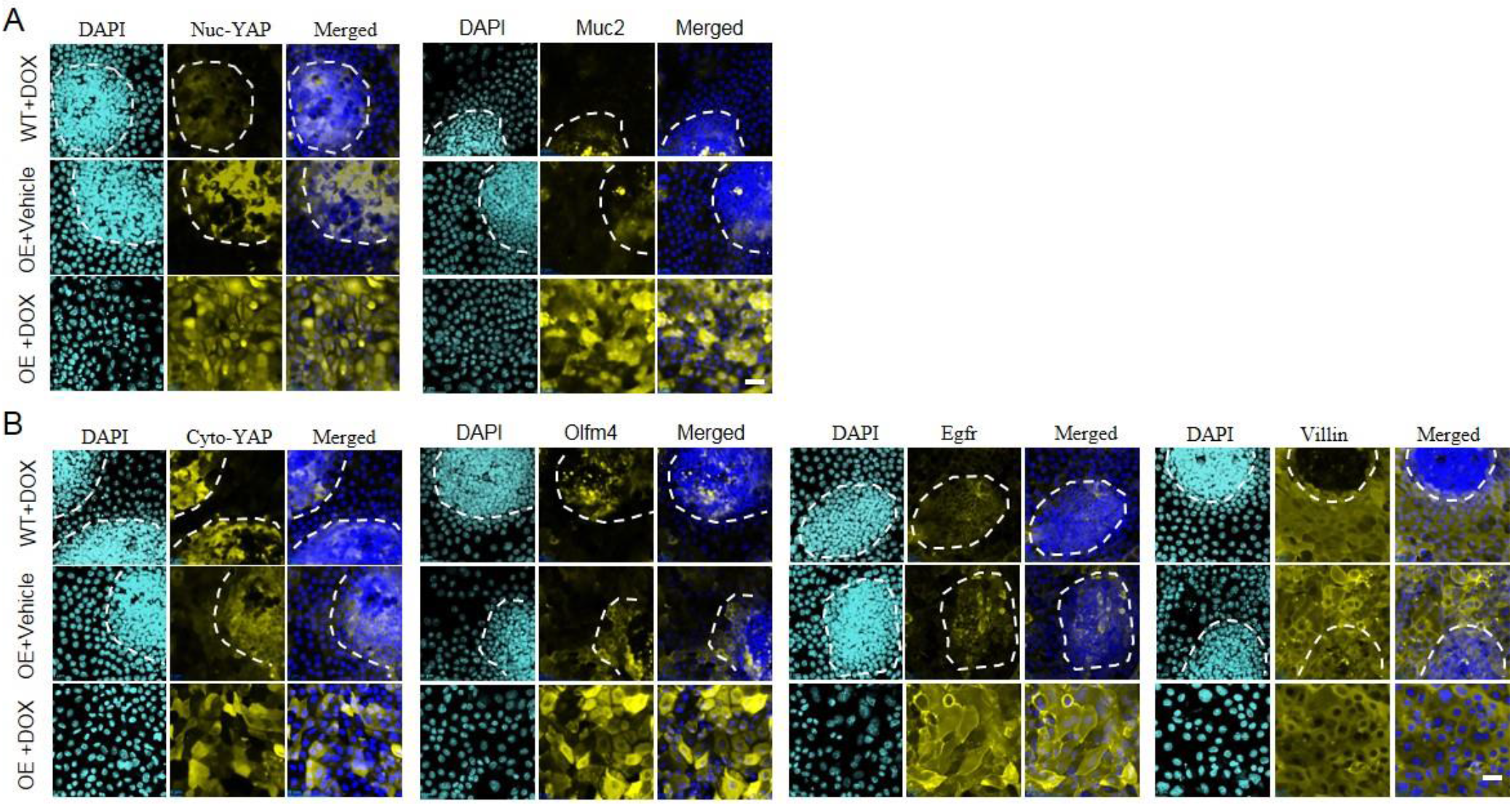
Staining for WT+DOX, YAP OE+Vehicle, and YAP OE+DOX. (A) Increasing nuc-YAP expression by OE promoted Muc2. (B) The increase in cyto-YAP augmented the expression of Olfm4 and Egfr. The white dashed lines trace the crypt-like regions. *n*=3. Scale bar, 25 μm.

**Extended Data Figure 14.**
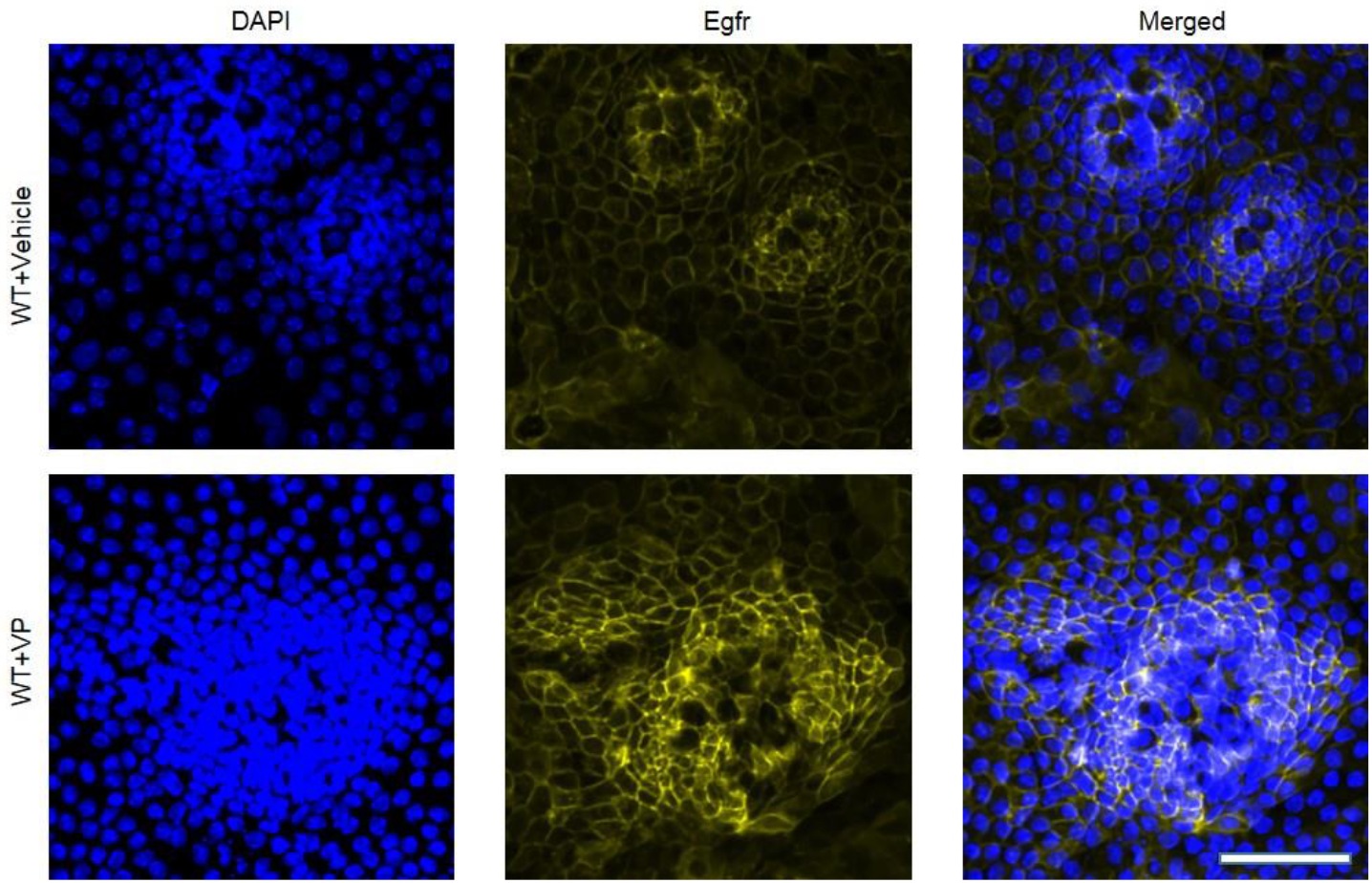
Staining for WT+Vehicle and WT+VP. VP augmented the expression of Egfr. *n*=3. Scale bar, 50 μm.

**Extended Data Figure 15.**
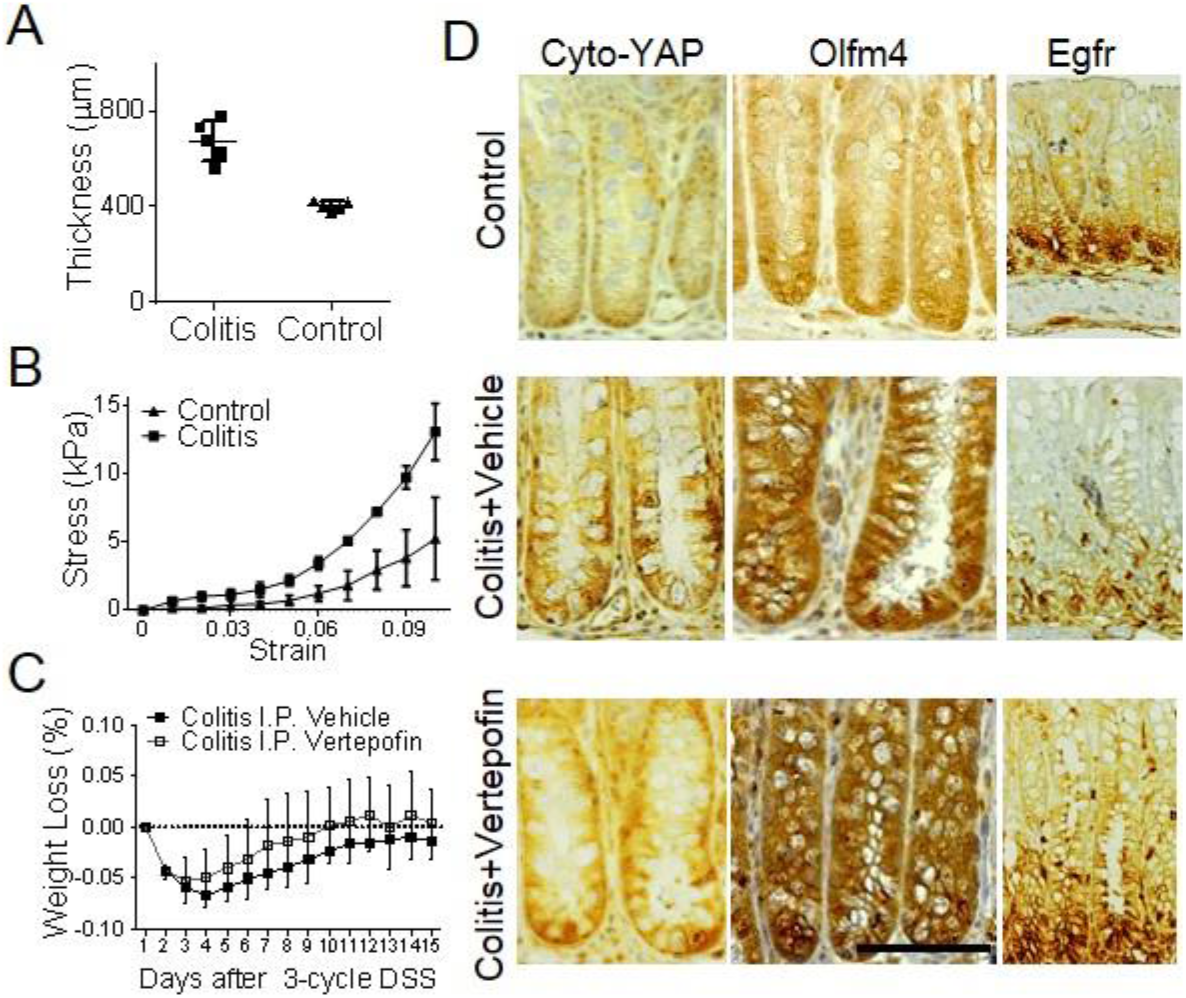
Colon thickened (A, *n*=6) and stiffened (B, *n*=3) in colitis group compared to the control. (C) VP treatment mitigated the body weight loss of the colitis group (*n*=6). (D) Cyto-YAP and Olfm4 were lower in control than the other two groups. VP treatment significantly increased the expression of Egfr. Scale bar, 200 μm.

**Extended Data Figure 16.**
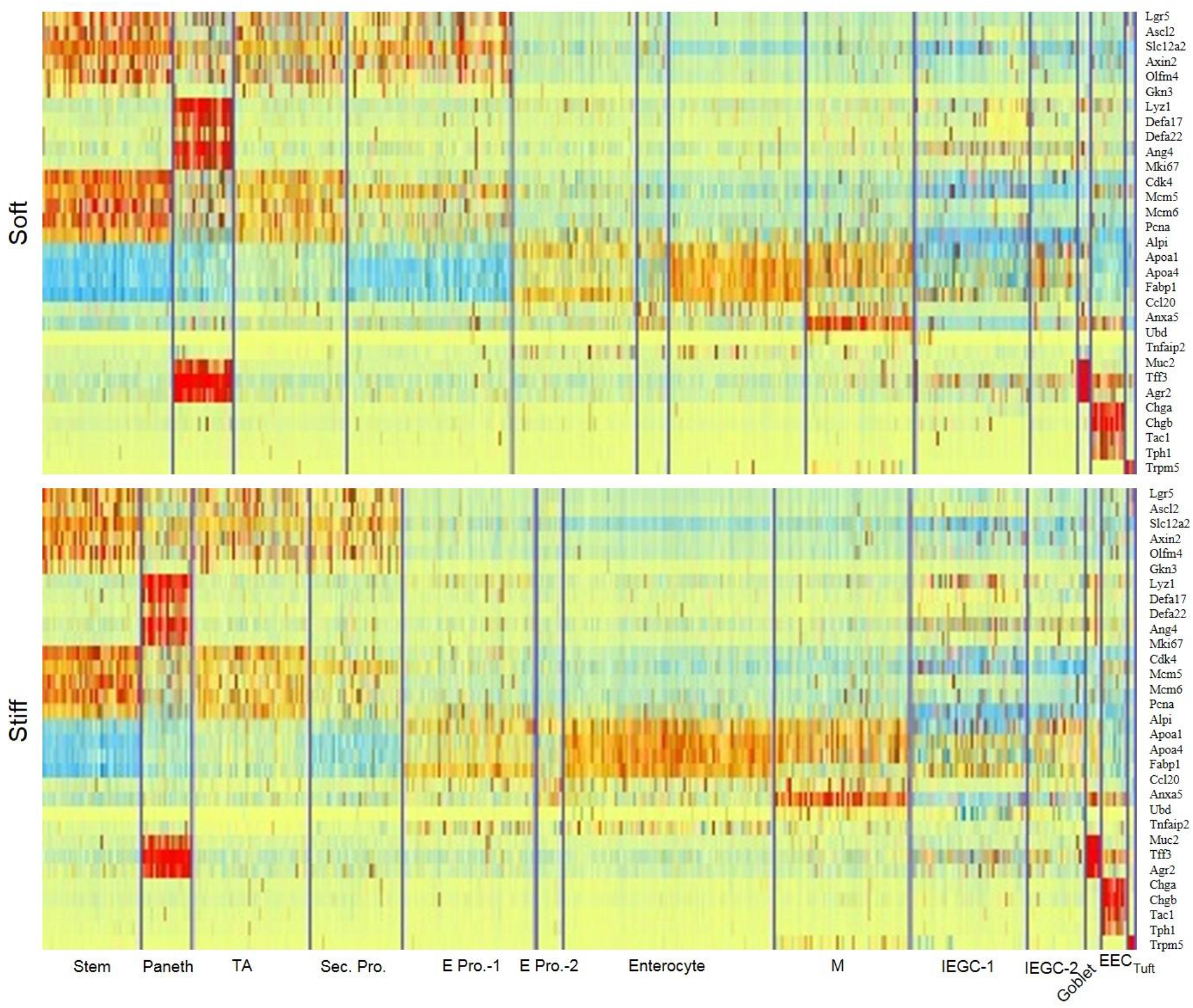
Full labels of marker genes for each cell type.

**Extended Data Figure 17.**
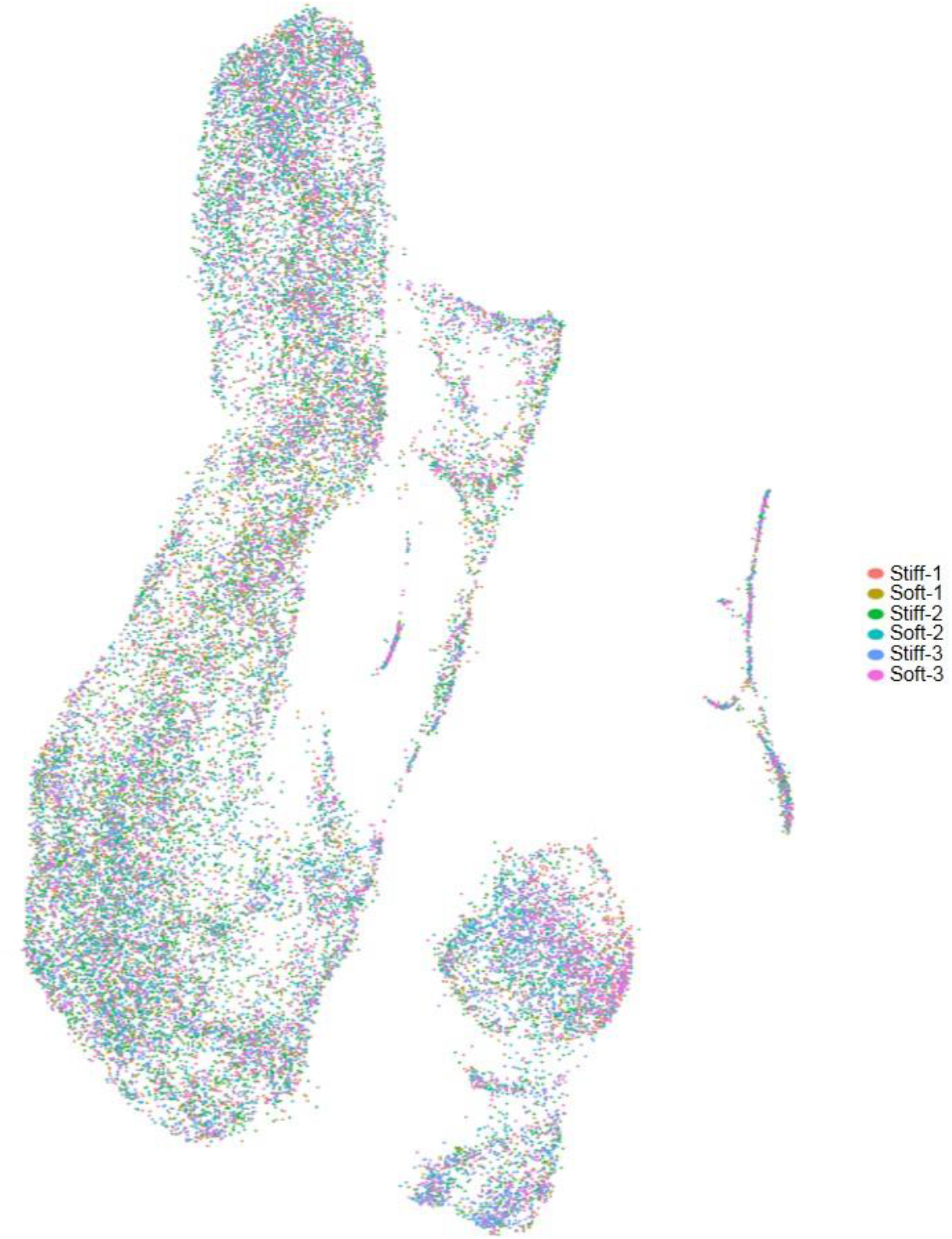
Three animals were used to triplicate the single-cell expression profiles. The clustering was consistent among the triplication on both the soft matrix and the stiff martix.

**Extended Data Figure 18.**
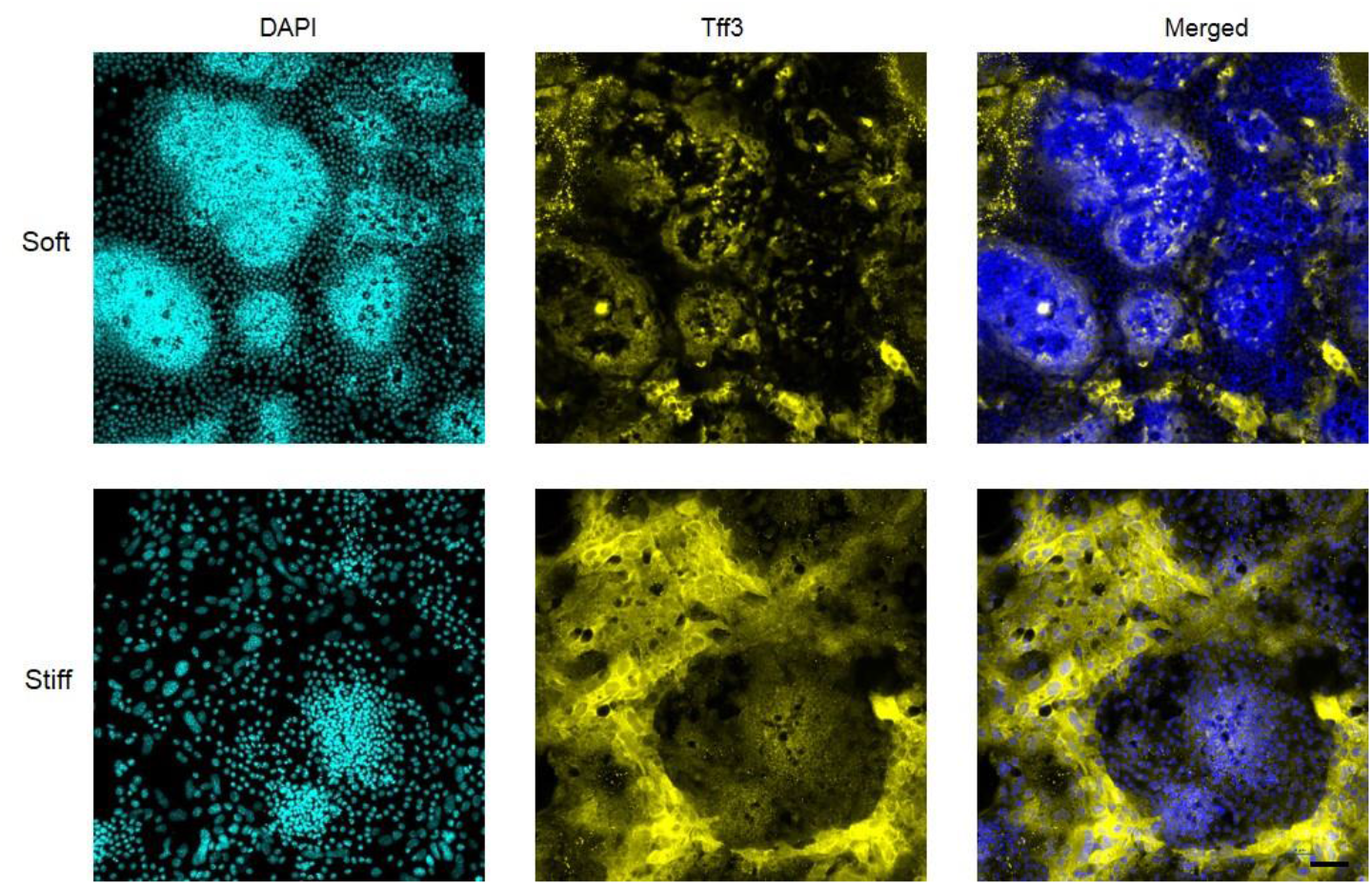
The goblet cell marker-Tff3 was increased on the stiff matrix (*n*=3). Scale bar, 25 μm.

**Extended Data Figure 19.**
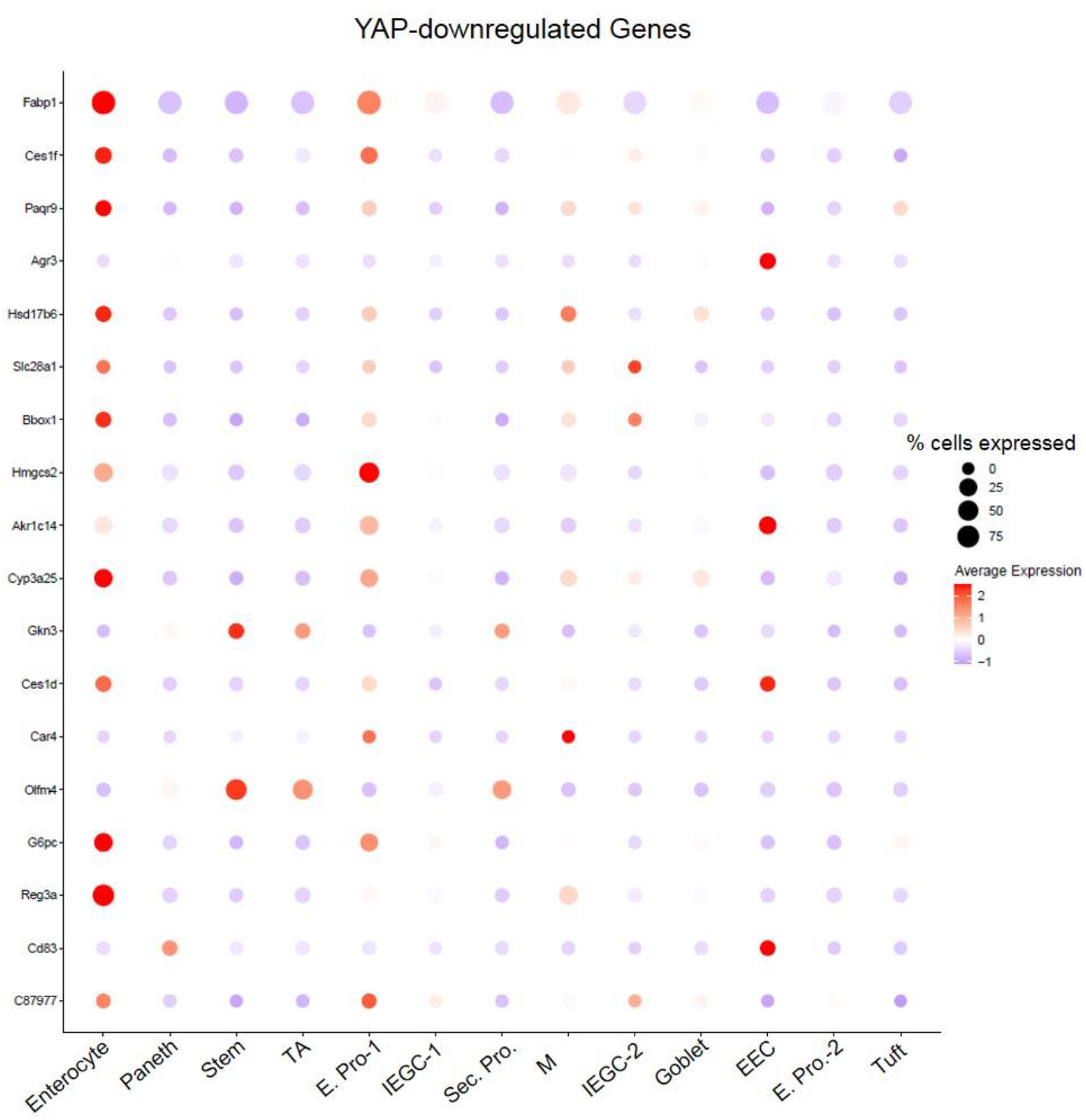
The genes downregulated by YAP highly expressed in enterocytes and their progenitors-1 (*n*=3).

**Extended Data Figure 20.**
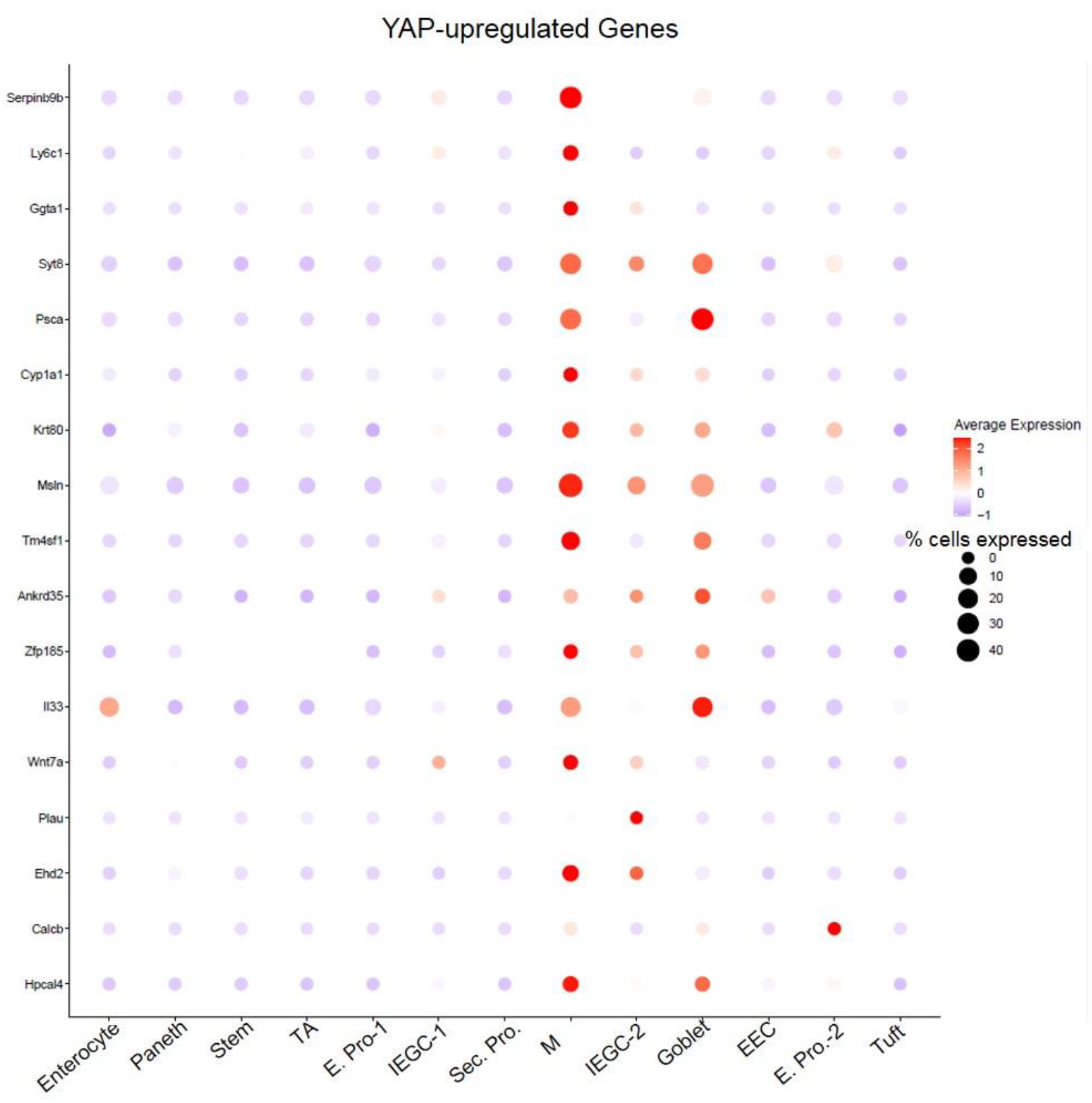
The genes upregulated by YAP highly expressed in goblet cells, IEGCs and M cells (*n*=3).

**Extended Data Figure 21.**
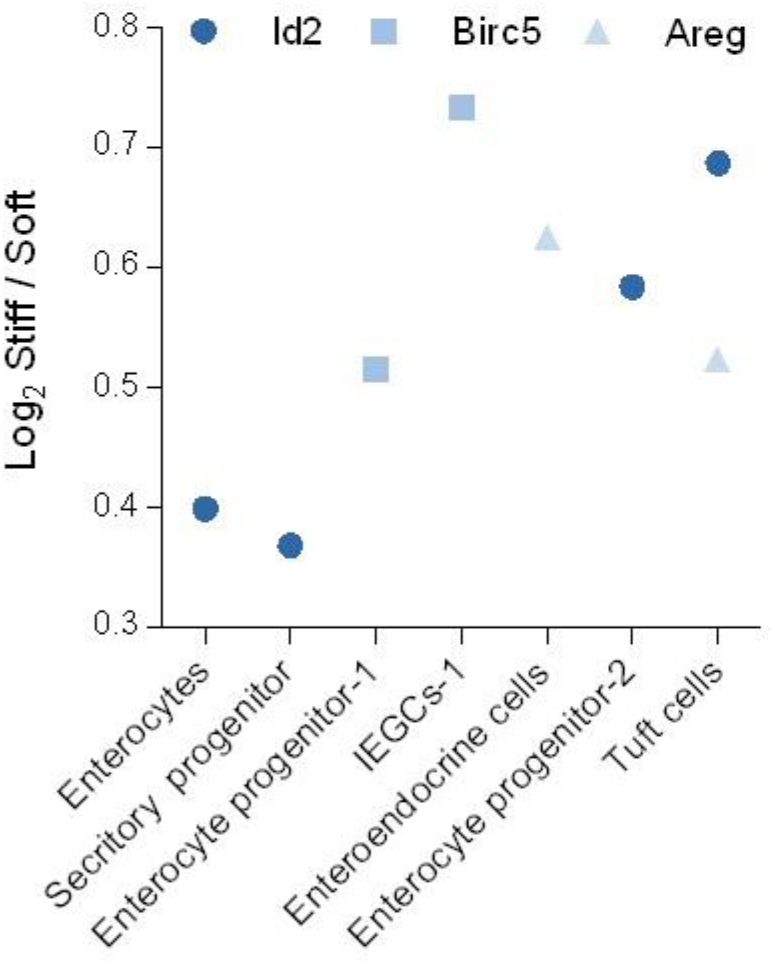
Differential gene expressions analysis shows the changes of downstream genes of nuc-YAP (*n*=3).

**Extended Data Figure 22.**
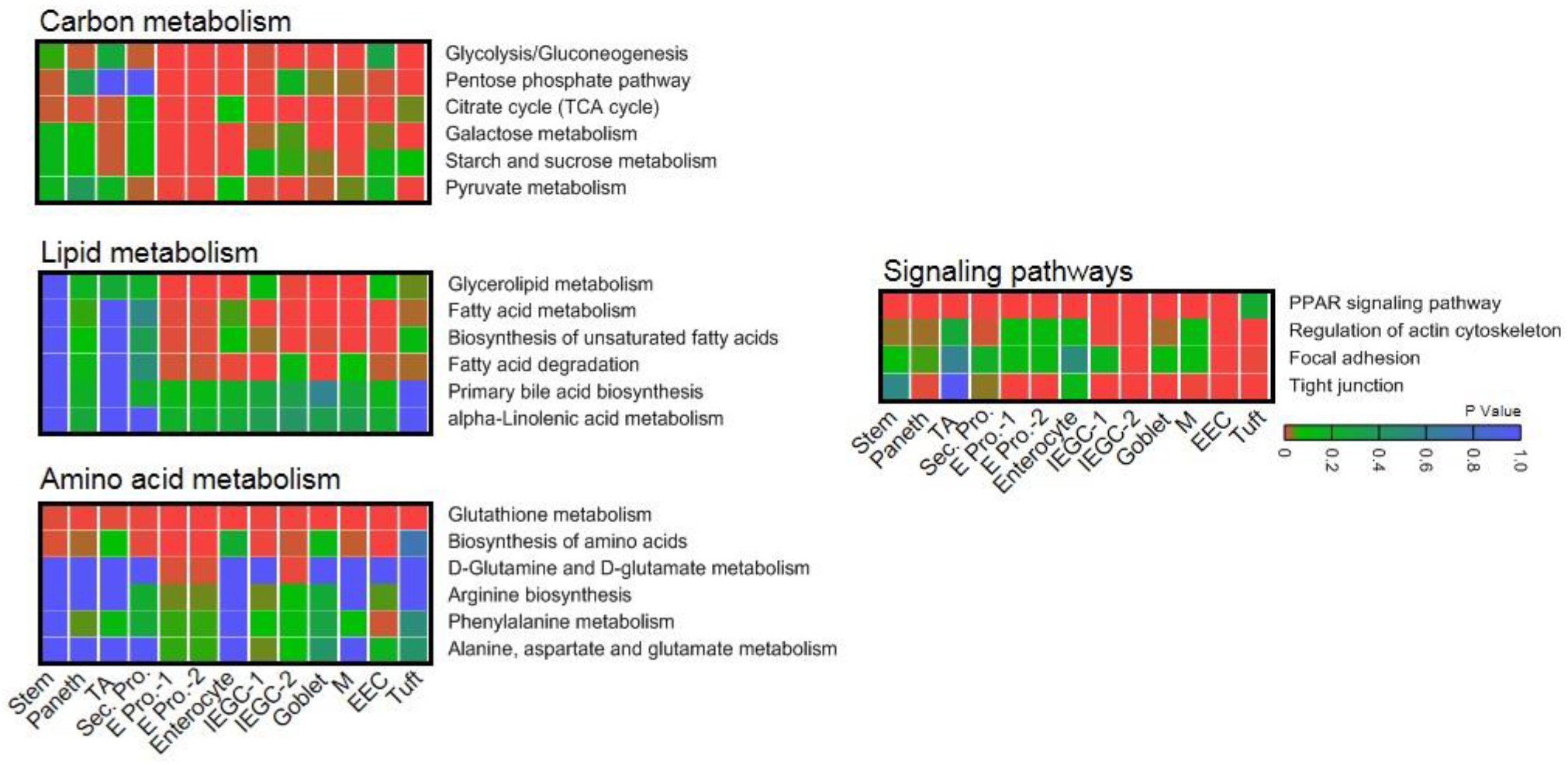
Pathway enrichment analysis is performed for carbon metabolism, lipid metabolism, amino acid metabolism and the signaling pathways. Compared to the soft matrix, carbon metabolism is more enriched than amino acid metabolism on the stiff matrix. Mechenotransduction-related signaling as well as Peroxisome proliferator-activated receptor (PPAR) was also more enriched on the stiff matrix (*n*=3).

**Extended Data Figure 23.**
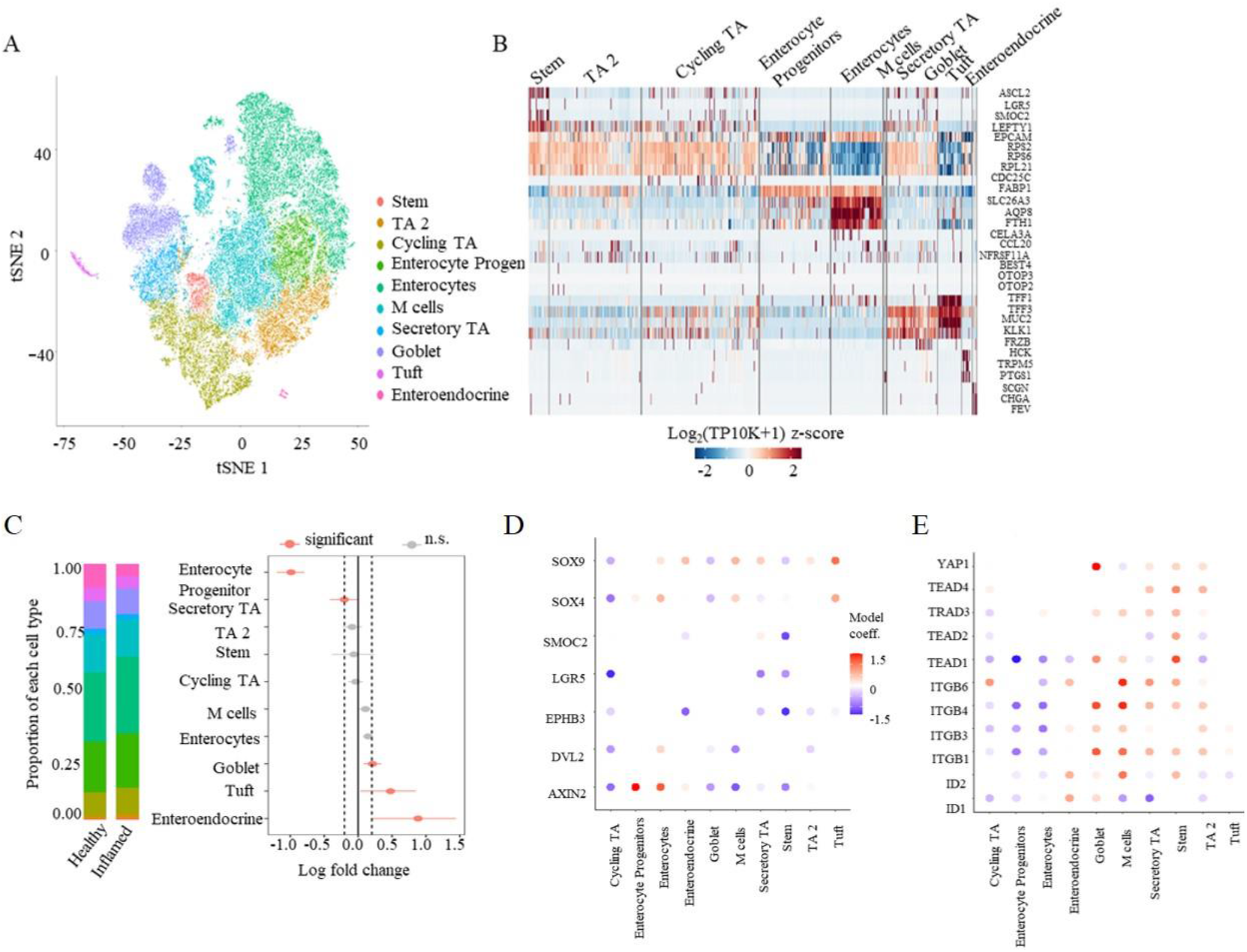
ScRNAseq analysis from healthy tissue and inflamed tissue biopsied from Ulcerative Colitis patients (*n*=3) and healthy individuals (*n*=5). (A) T-Stochastic Neighborhood Embedding (t-SNE) of cells colored by cell type from all samples. (B) Marker genes for each cell types. (C) Left panel: Average cell type proportions in aggregates of healthy or inflamed samples. Right panel: Fold changes in proportion of each cell type in UC patients compared to healthy individuals. Whiskers correspond to highest and lowest points within 1.5 interquartile range. Significant criteria, *P*<0.05. (D) The Wnt pathways (e.g. Sox4, Sox9, Lgr5) suppressed specifically by YAP are downregulated in the ISCs of UC. Model coefficient reported here is the discrete component of the hurdle model. (E) The mechanosignaling pathway including integrin (ITGB), YAP, and TEAD is highly activated in both ISCs and the secretive cell types of UC, but not in enterocytes.

**Extended Data Figure 24.**
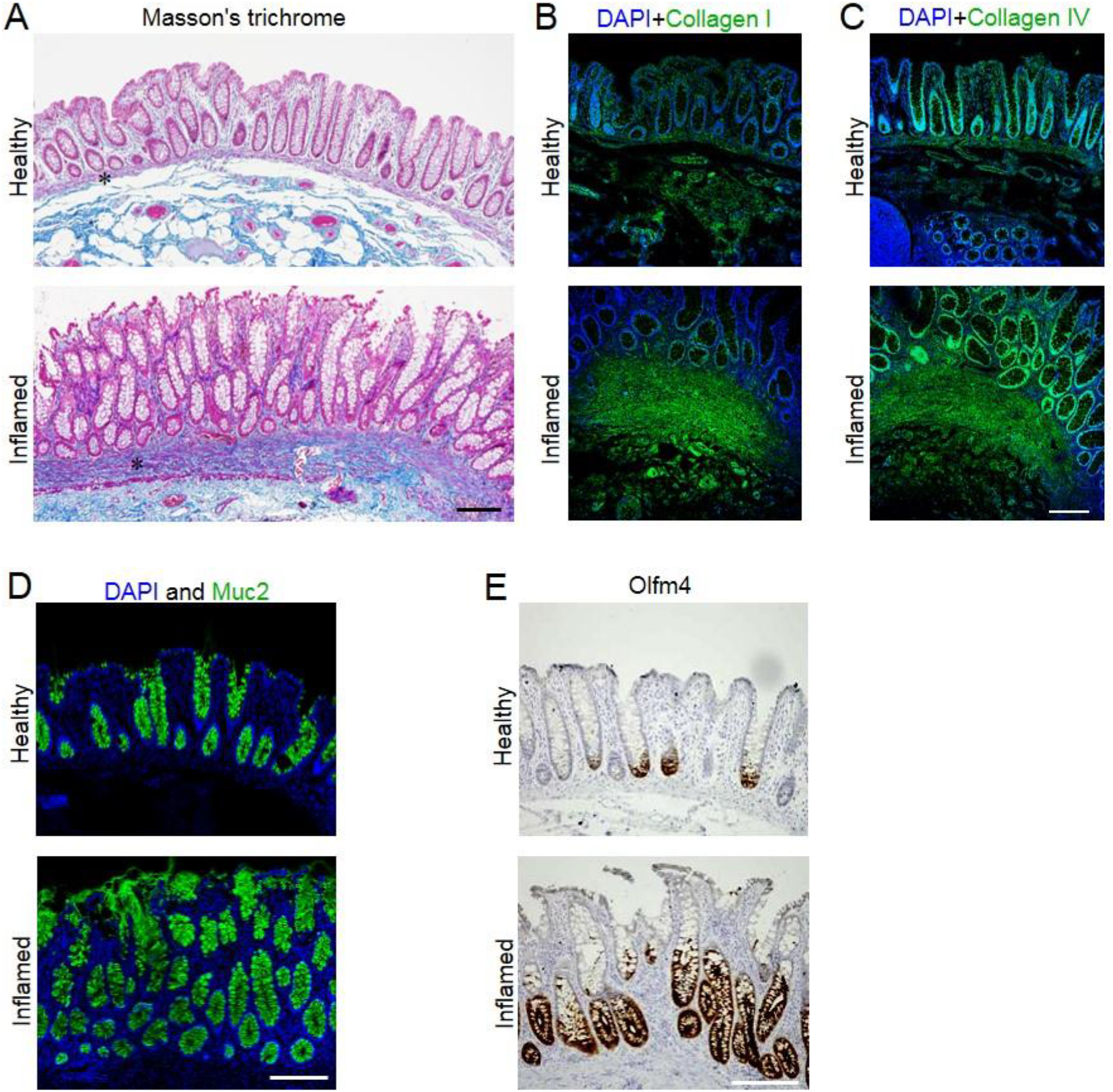
The masson’s trichorme staining (A) and the staining of collagen I (B) and Collagen IV (C) showed the fibrosis and thickening of BM and lamina propria labelled with asterisks. (D) Muc2^+^ goblet cells increased in the inflamed colon. (E) Olfm4 was increased and expanded into larger regions in the inflamed colon. *n*=3. Scale bar, 200 µm.

**Extended Data Figure 25.**
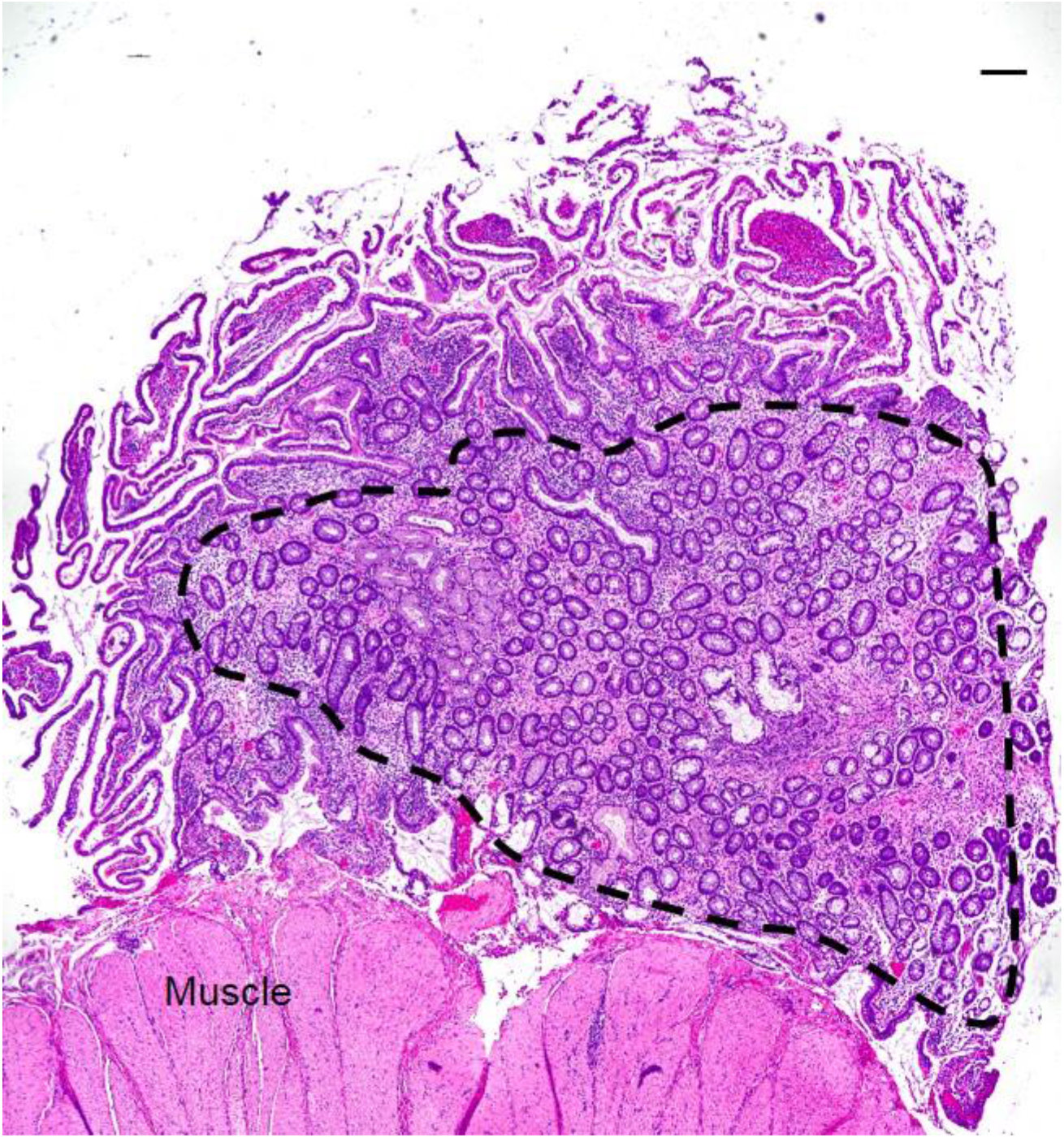
In a large area (outside of the dashed line) of structured ileum (extreme fibrosis), the invaginated ISC niche-crypts nearly disappeared and only pieces of the villi were left, resembling the stiffness-reduced size of the crypt and loss of ISCs. Meanwhile, lots of ectopic crypts (inside the dashed line) formed, resembling stiffness-induced new crypt formation. *n*=1. Scale bar, 200 µm.

**Extended Data Figure 26.**
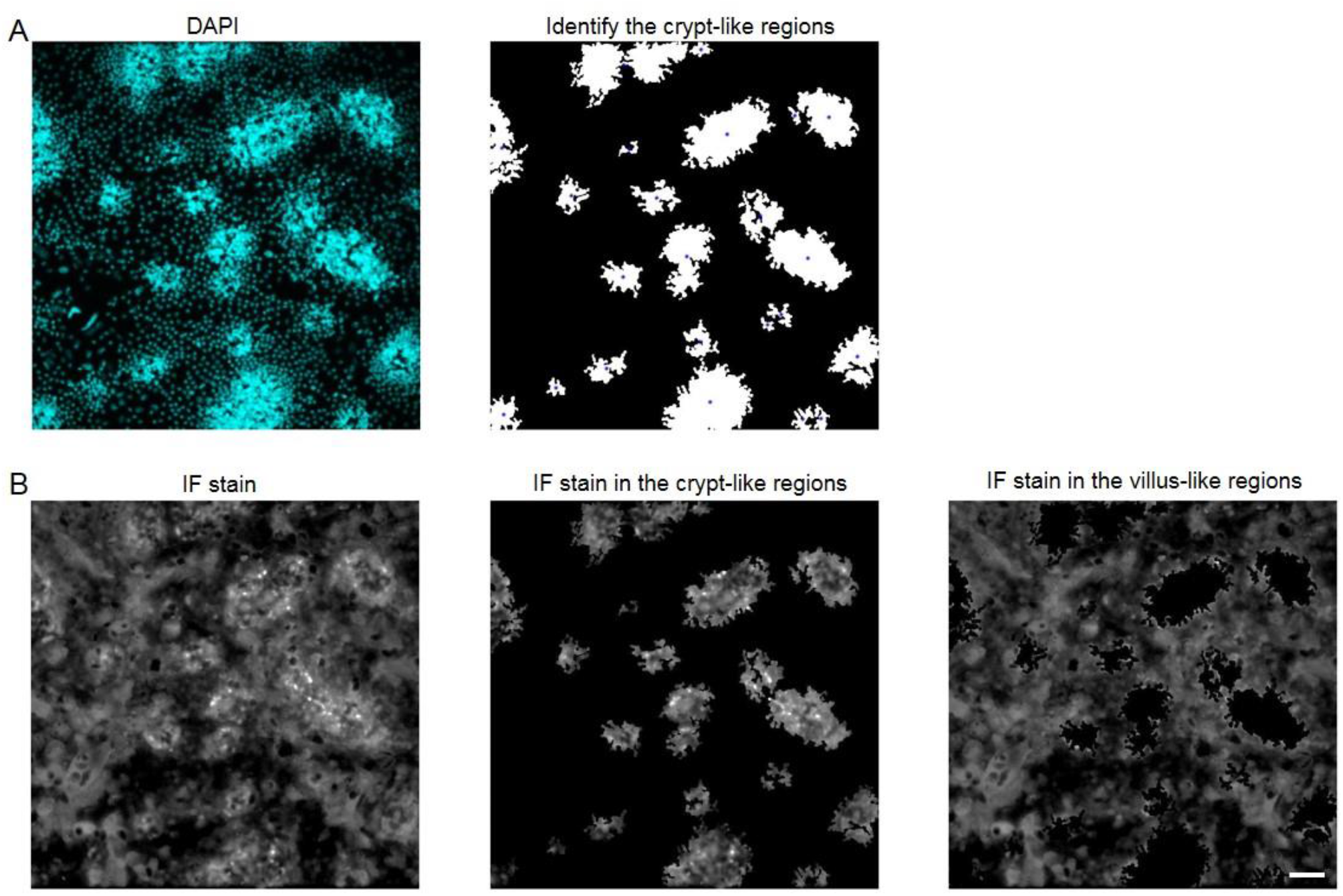
Illustration for the customized MATLAB code. (A) The crypt-like regions were identified based on the intensity of the DAPI staining. (B) The fluorescent signals were respectively isolated in the crypt-like regions and the villus-like regions. Scale bar, 100 µm.

**Extended Data Figure 27.**
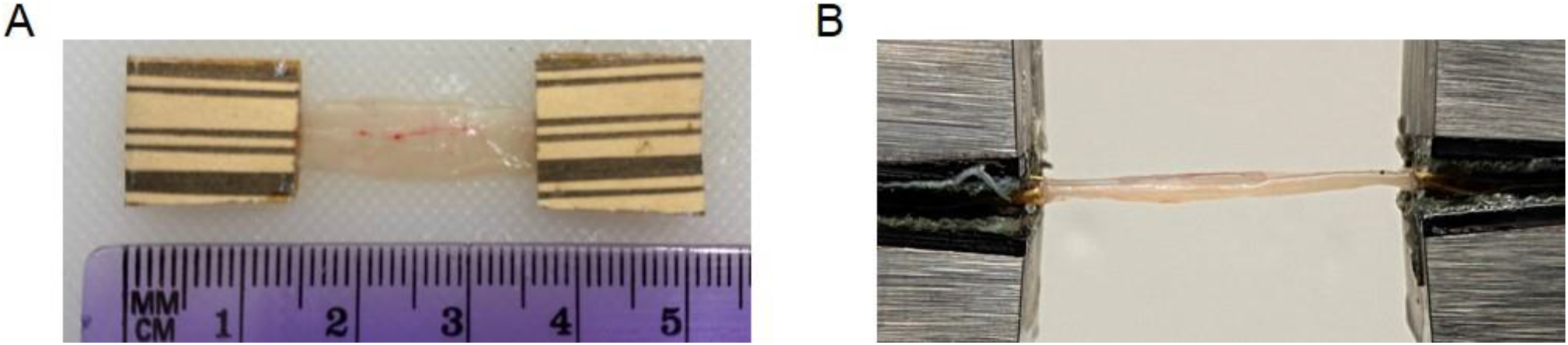
(A) An intestinal tissue sample with sandpaper tabs at both ends; and (B) uniaxial tensile test of the intestinal tissue sample.

